# Transcription factor collaboration enables precise T cell state engineering

**DOI:** 10.64898/2026.04.20.718569

**Authors:** Rachel E. Savage, Christian D. McRoberts Amador, Conrad T. Hock, Ruochi Zhang, Hung-Che Kuo, Aretha R. Gao, Max A. Horlbeck, Charles A. Gersbach, Jason D. Buenrostro

## Abstract

Transcription factors (TFs) collaborate to regulate gene expression programs that define cell fate. In CD8^+^ T cells, this coordinated regulation underlies exhaustion, a dysfunctional state that constrains immunity in chronic infection and cancer. Here, we screen for cell state-specific TFs by performing pooled overexpression screens of 3,548 TF and TF isoforms in primary T cells across multiple CD8^+^ T cell states. We identify 82 regulators that collaborate with exhaustion-specific programs and profile their effects using perturb-SHARE-seq, connecting perturbations to changes in chromatin accessibility and gene expression across 702,314 single cells. We identify 38 reproducible regulatory programs and construct a map of 12,616 TF-program connections that shape CD8^+^ T cell states, nominating KLF2 as predictive of positive response to CAR-T therapy. Using seq2PRINT, a deep learning framework that predicts functional TF interactions, we identify RUNX as a “master collaborator”, a TF that broadly collaborates with other factors, and uncover a RUNX2:KLF2 interaction that specifies exhaustion-associated programs. Mutation of the RUNX2:KLF2 protein interface attenuates KLF2-mediated repression of exhaustion, while synthetic tethering of RUNX2 to KLF2 leads to an amplification of the phenotype. More broadly, we identify the collaborative action of RUNX as a driver in CD8^+^ T cell states, and show that tethering TFs enables the rational engineering of cell state identity for cell and gene therapies.

## Introduction

Cells exist in a diverse range of types and states, each enabling specialized functions within tissues and dynamic adaptations to environmental cues^1^. This diversity is molecularly encoded in the binding of transcription factors (TFs) across the genome that establish and maintain distinct gene expression programs^2,3^. TFs are collaborative, co-binding with multiple TFs to regulate gene expression^4^. Single-cell genomics and large-scale functional screens have enabled the discovery of TFs associated with diverse cell states^5–10^. Taken together, these advances have made cell state engineering increasingly feasible^8,11–13^. However, reprogramming cell identity often requires two or more TFs, and the discovery of these combinations represents a field-defining challenge that precludes the rational design of cell types^14–16^. Here, we hypothesize that a molecular understanding and targeting of TF interactions will enable new opportunities to rationally design cell identity.

The design gap is particularly evident in CD8^+^ T cells, whose differentiation into memory, effector, and exhausted states critically shapes infection, cancer, and immunotherapy outcomes^17^. Single-cell atlases have defined diverse CD8^+^ T cell states across health and disease^18^ and gain- and loss-of-function studies have nominated factors capable of shifting cells between these states^8–10,19–23^. However, while T cells can be engineered to target tumor cells^24^, the clinical success of immunotherapy has been limited by an incomplete ability to prevent or reverse exhausted T cell states, a state characterized by sustained inhibitory receptor expression and a diminished capacity to mount cytotoxic responses^17^. We hypothesized that TF collaboration could be leveraged to more effectively reprogram CD8^+^ T cell states. However, TF interactions span a vast combinatorial space, rendering systematic interrogation of collaboration largely intractable.

Here, we performed a pooled overexpression screen to identify context-specific regulators of CD8^+^ T cell exhaustion. To enable mechanistic dissection of the resulting regulators, we performed perturb-SHARE-seq for simultaneous measurement of single-cell gene expression, chromatin accessibility, and perturbation identity. Across 702,314 cells, we map the effects of key regulators on cell identity and identify composite regulatory elements as defining features of cell state. We identify RUNX as a master collaborator, a central hub that collaborates with diverse TFs, and find that composite RUNX motifs are more state-specific than monomers, indicating that collaborative TF binding, rather than single TF binding, delineates cell states. Focusing on exhaustion-associated programs, we identify cooperative interactions between RUNX2 and KLF2 at key exhaustion-related loci. Mutagenesis demonstrates that RUNX2 and KLF2 cooperation modulates repression of exhaustion programs. Synthetic tethering of RUNX2 and KLF2 is sufficient to amplify this regulatory effect and more effectively reprogram T cell state than either factor alone or the untethered counterparts. Together, these findings establish TF collaboration at composite regulatory elements as a mechanistic unit of CD8^+^ T cell state specification and provide a framework for rational engineering of immune cell identity.

## Results

### A TF-wide gain-of-function screen reveals regulators of TOX

We sought to identify TFs that regulate CD8^+^ T cells specifically in the context of exhaustion. This approach enriches for factors that modulate endogenous exhaustion programs without broadly reprogramming cell identity, and for those that may act in concert with endogenous cofactors uniquely available in the exhausted state. To identify such regulators, we assessed the ability of TFs to alter levels of TOX, a key driver of T cell exhaustion^25^, across multiple CD8^+^ T cell states. We performed pooled gain-of-function screens using the MORF library, which encodes 3,548 TFs and TF isoforms^12^. Primary CD8^+^ T cells from three human donors were transduced with the MORF library, then cultured under standard expansion conditions (“expanded”), or under chronic α-CD3/CD28 stimulation conditions (“exhausted”), a well-established *in vitro* model of exhaustion^9,26^ (**Fig. 1A**). Finally, we used FACS to isolate cells with high (TOX^HIGH^) or low (TOX^LOW^) TOX protein levels (**Fig. 1A, Extended Data Fig. 1**).

**Figure 1.**
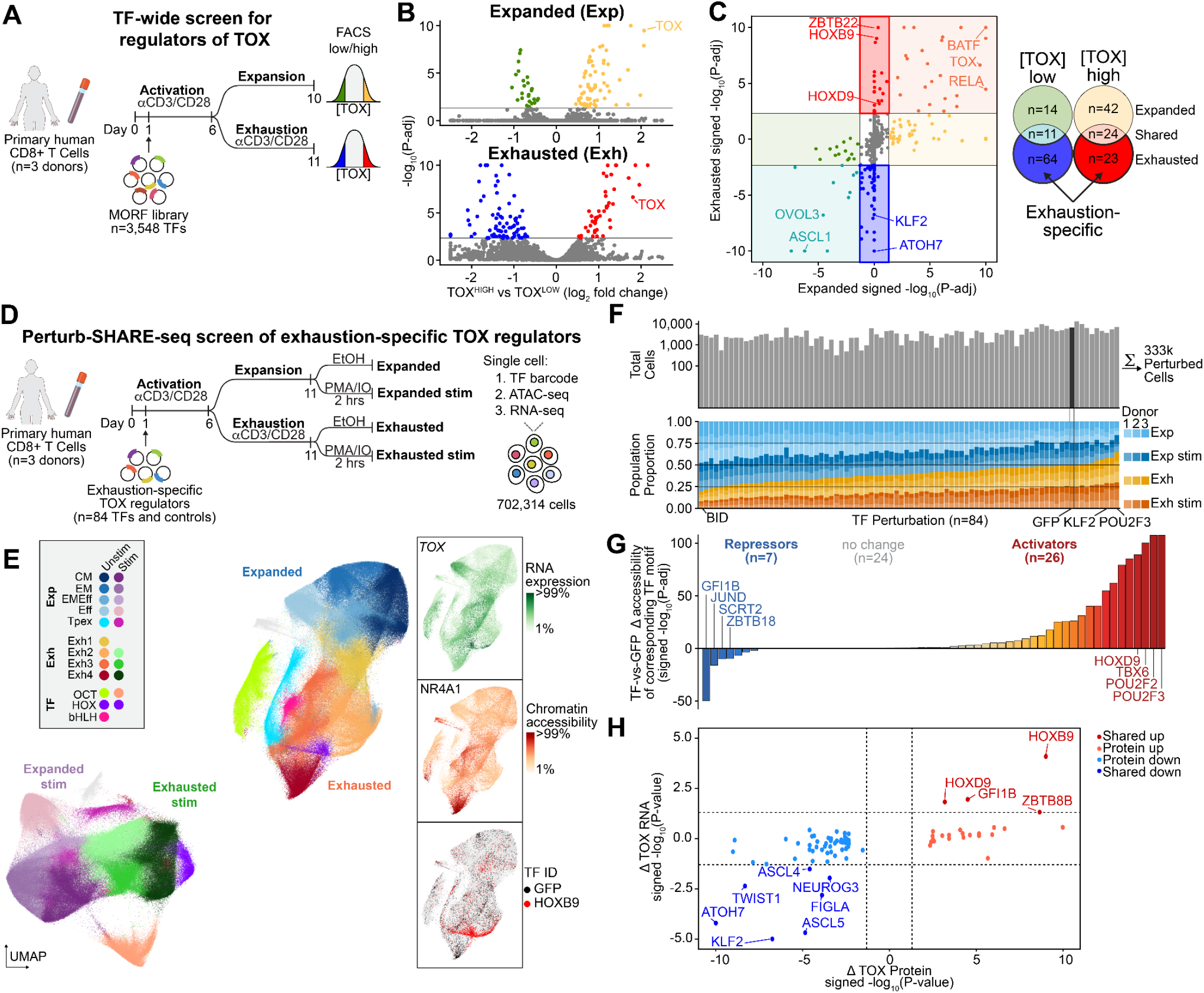
Identification and characterization of master regulators of CD8^+^ T cell states. (A) Schematic of TF-wide screen for regulators of TOX in primary human CD8^+^ T cells. (B) Volcano plots of differential enrichment of TF barcodes in the TOX^HIGH^ versus TOX^LOW^ populations in expanded (top) and exhausted (bottom) conditions. (C) Comparison of signed –log_10_ adjusted *P* values in expanded (x-axis) versus exhausted (y-axis) conditions. TFs that uniquely decrease (blue) or increase (red) TOX expression in the exhausted condition are highlighted. (D) Schematic of perturb-SHARE-seq experimental layout across four conditions: expanded, expanded stimulated, exhausted, and exhausted stimulated. (E) UMAP embedding of the perturb-SHARE-seq dataset (702,314 cells), highlighting TOX RNA expression, NR4A1 motif chromatin accessibility, and HOXB9 (red) and GFP (black) perturbed cells. (F) Recovery of cells expressing individual TFs across conditions, shown as absolute counts (top), and proportions (bottom) (333k cells). (G) TF-induced differential accessibility at each TF’s annotated motif, quantified as signed −log_10_ adjusted *P* values comparing TF-overexpressing cells to GFP controls. Labels denote activating (increased accessibility) and repressive (decreased accessibility) effects. (H) Comparison of TF-induced change in relative TOX protein levels (TF screen TOX^HIGH^ vs TOX^LOW^ signed -log_10_ *P*) versus TOX RNA levels (perturb-SHARE-seq TF vs GFP signed -log_10_ *P*).

Across both conditions, we identified 133 TFs that increased and 172 that decreased TOX expression (p-adj<0.05, |log_2_FC|>0.5) (**Fig. 1B, Extended Table 1**). As expected, TOX overexpression itself produced robust increases in TOX protein under both expanded (log_2_FC=2.07, p-adj=3.311x10^-10^) and exhausted (log_2_FC=1.8, p-adj=2.30x10^-7^) conditions. Additionally, across both conditions, we recovered 21 shared positive regulators, including factors linked to BAF/AP-1 complexes (BATF)^27^, NF-κB signaling (RELA)^28^, ER stress responses (XBP1)^29^, and Myc-driven activation (MYC, MYCL)^30^, all pathways previously implicated in T cell activation and exhaustion. We discovered 17 TFs that decreased TOX protein levels in both conditions. Several hits correspond to lineage-specifying TFs, including epithelial (OVOL1/3), mature neuronal (ASCL1/3, NEUROD4/G1), and NK-like (NFIL3) lineages. The recovery of established TOX regulators together with previously uncharacterized modulators underscores the breadth of pathways influencing TOX abundance.

We identified 87 TFs that uniquely increased (n=23) or decreased (n=64) TOX in the exhausted condition (**Fig. 1C, Extended Table 1**). Many of the 23 TFs, or their paralogues, that upregulated TOX have been implicated in CD8^+^ differentiation and dysfunction, such as BATF2^20^, NFKBID^31^, POU2F2-3^18^, ID1-3^32^, and HOXB9/D9^33^. We also identified novel positive regulators, including zinc finger proteins (GFI1B, ZBTB18, ZBTB22/34/8B, ZNF575), pioneer-like TFs (LHX6, PRRX1, MEOX1, FOXH1), and epigenetic regulators (WHSC1, EWSR1). Among the 64 TFs that decreased TOX in exhaustion, we observed enrichment for neurogenesis-associated TFs (n=25, GO FDR=1.617x10^-4^), including bHLHs (ASCL2, ATOH7, NEUROG3, TCF3, TWIST1) and homeoboxes (BARHL2, DBX1, EN1-2, GBX2, GSX2, IRX5, PTF1A, MNX1). This may reflect neurogenic regulation of TOX during corticogenesis^34^. Consistent with prior literature^35^, we also found KLF2 decreased TOX levels (log_2_FC=-1.72, p-adj=1.78x10^-7^). Overall, the screen identified both previously reported and uncharacterized regulators.

### Single-cell multiomic screening of exhaustion-unique TF perturbations

We sought to elucidate the mechanisms by which TFs altered TOX expression in exhausted CD8^+^ T cells, therefore we profiled 82 of the exhaustion-unique hits. Resolving these regulatory effects at the resolution of chromatin engagement of TFs requires joint measurement of chromatin accessibility, gene expression, and perturbation identity in single cells. Existing single-cell perturbation multi-omic approaches are limited by cost and scale. We therefore developed perturb-SHARE-seq, an extension of SHARE-seq^36^ that captures single cell perturbation barcodes alongside gene expression and chromatin accessibility profiles and incorporates an improved computational framework for perturbation assignment (**Extended Data Fig. 2**, see Methods).

We performed perturb-SHARE-seq on 82 exhaustion-specific TFs together with TOX and GFP controls (n=84 TFs and controls) across three human donors. CD8^+^ T cells were transduced with the pooled TF library, selected with puromycin, and cultured for 11 days under expansion or exhaustion conditions. Both the expanded and exhausted conditions were further split into phorbol 12-myristate 13-acetate and ionomycin (PMA/IO) stimulation and control conditions, resulting in four distinct conditions (**Fig. 1D**). We recovered high-quality single cell libraries, with median library sizes of 3,893 and 2,784 reads per cell for ATAC and RNA, respectively, yielding a final dataset representing 702,314 single cells including 333,035 cells with assigned perturbation labels (47% of all cells, FDR < 0.05) (**Fig. 1E, F** and **Extended Data Fig. 2**). In total, this dataset comprised 1,992 measurement profiles (83 perturbations x 3 donors x 4 conditions x 2 modalities).

Cells largely clustered according to known CD8^+^ state markers, with the majority of clustering driven by stimulation and exhaustion state (**Fig. 1E, Extended Data Fig. 3**). Exhausted cells displayed increased expression of exhaustion-associated genes such as *TOX* and *CTLA4*, increased chromatin accessibility at exhaustion-associated TF motifs such as NR4A1^37^, and decreased expression of activation genes such as *IFNG* following PMA/IO stimulation (**Fig. 1E, Extended Data Fig. 3**). Memory and effector cells were marked by *BCL2* and *GZMB*, respectively. Additionally, we found clusters (n=4) that were associated with TF perturbations, including two OCT-specific clusters (POU2F2, POU2F2 isoform 2, POU2F3), a HOX-specific cluster (HOXB9, HOXD9) and a cluster enriched for bHLH TFs (ASCL2,4,5; ATOH7; BHLHA9; FIGLA; MSC; NEUROG3; PTF1A; TWIST1) (**Extended Data Fig. 4**).

To assess whether perturb-SHARE-seq captures perturbation-driven effects on gene regulation, we examined TF-driven changes in chromatin accessibility and gene expression. We hypothesized that overexpressed TFs preferentially alter chromatin accessibility at the TF’s specific binding motif (i.e., cells overexpressing POU2F3 would have increased accessibility of POU2F3 motifs). Indeed, 33 of 57 TFs that had annotated motifs showed significant changes in chromatin accessibility at their motifs (p-adj < 0.05), with activators (n=26) increasing and repressors (n=7) decreasing accessibility (**Fig. 1G, Extended Data Fig. 2**). Repressive TFs such as GFI1B and SCRT2, which endogenously recruit LSD1 to silence chromatin^38^, recapitulated their known repressive function, indicating that exogenous TF expression mimics endogenous regulatory activity. We additionally observed strong concordance between TF effects on TOX RNA levels, as measured in perturb-SHARE-seq, and relative TOX protein levels, as measured by the FACS-based screen, with 69 of 83 TFs showing consistent directional effects (**Fig. 1H**). Together, these results establish perturb-SHARE-seq as a scalable, mechanistically informative readout of TF-driven chromatin and transcriptional regulation.

### TF perturbations modulate clinically relevant chromatin and gene expression programs

TFs directly and indirectly regulate gene expression programs to alter cell states. To capture this regulatory structure, we decomposed single-cell chromatin accessibility and RNA expression into robust programs. Briefly, we adopted an iterative subsampling framework^39^, and retained topics derived from cisTopic^40^ Latent Dirichlet Allocation (LDA) modeling that were consistent across multiple iterations of subsampling (**Fig. 2A**). Altogether, we identify 38 reproducible ATAC (n=21) and RNA (n=17) programs (**Fig. 2B** and **Extended Data Fig. 5**). Overall, ATAC and RNA programs captured a range of CD8^+^ T cell states and functions, ranging from memory, effector, exhausted, precursor exhaustion, and cell cycle programs (**Extended Data Fig. 5**). Single cell program scores were state specific, and program gene markers overlapped with known T cell gene sets (**Extended Data Fig. 3C, 5**). To highlight one example, ATAC program 1 and RNA program 1 mark the same population of cells (**Fig. 2C**), corresponding to an effector memory-like state, enriched for canonical markers such as *IL7R* and memory signatures^41^. Together, these programs capture coherent and reproducible cell states, providing a framework for linking TF activity to cell identity.

**Figure 2.**
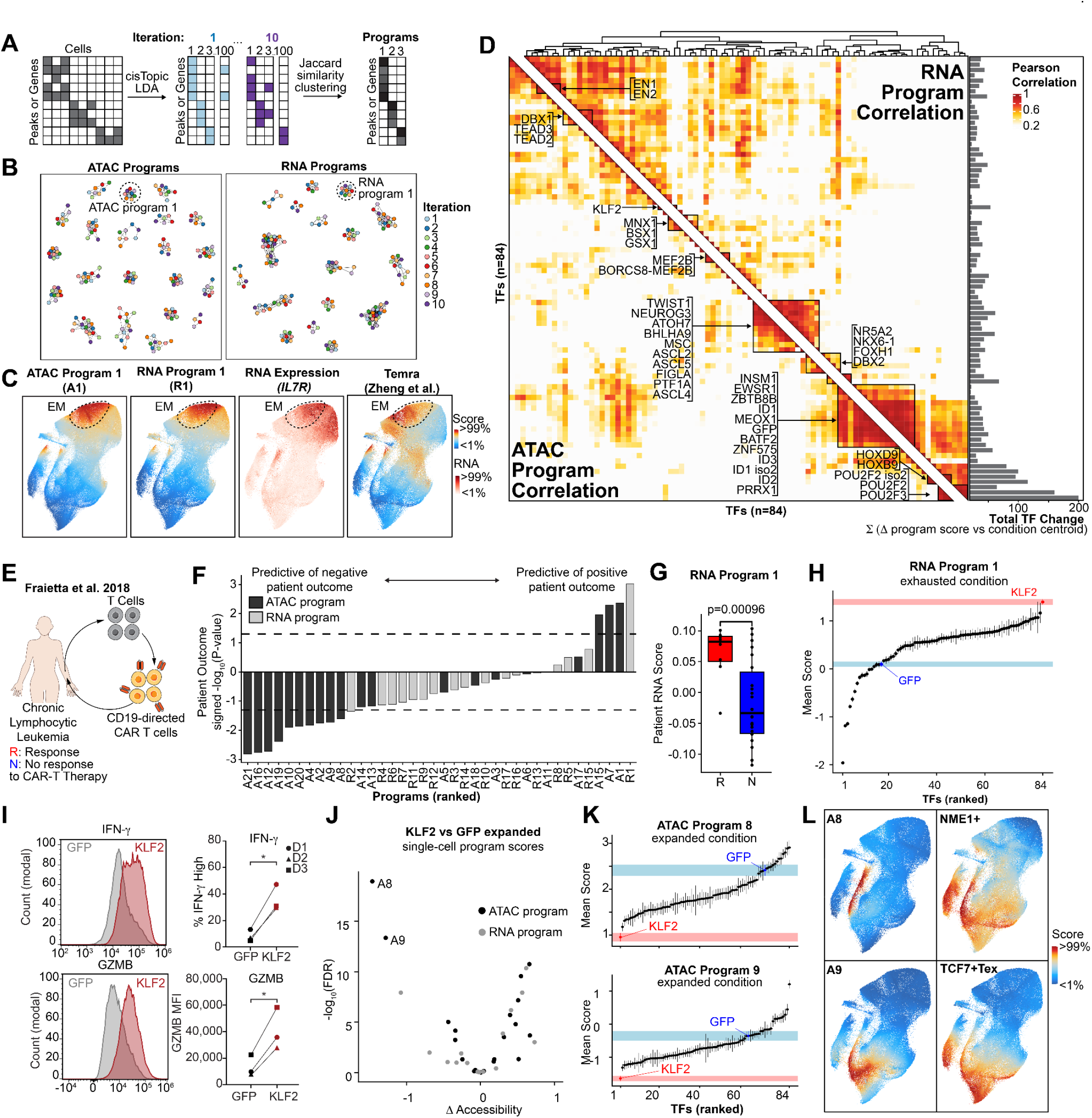
Transcription factor perturbations reshape regulatory programs linked to therapeutic T cell response. (A) Schematic of the workflow to derive robust RNA and ATAC programs. Gene-by-cell (RNA) and peak-by-cell (ATAC) matrices were subjected to iterative topic modeling, retaining reproducible programs across runs. (B) Jaccard similarity networks of ATAC (left) and RNA (right) programs across iterations. Nodes represent topics (colored by iteration); edges denote overlap in peaks or genes. (C) UMAP of ATAC and RNA program 1 scores, overlapping the effector memory cluster (EM, dashed outline), and *IL7R* expression and Temra score^41^. (D) Heatmap of pairwise Pearson correlations between TF perturbations, partitioned by TF effects on ATAC programs (bottom left) and RNA programs (top right). Bar plot shows total perturbation-induced change. (E) Strategy for scoring patient bulk RNA-seq^42^ using RNA and ATAC programs. (F) Predictive significance of RNA (grey) and ATAC (black) programs for patient response, with responders assigned positive and non-responders assigned negative -log_10_ *P* values. (G) RNA program 1 score in CAR-T therapy responders (R) and non-responders (NR). (H) Expression of RNA Program 1 in exhausted cells across TF perturbations. Points indicate mean ± s.e.m.; GFP (blue) and KLF2 (red) highlighted. (I) Representative flow cytometry plots of Granzyme B and IFN-γ in exhausted cells following PMA/IO stimulation, comparing GFP (grey) and KLF2 (red) overexpression, with quantification (right, n=3). \**P* < 0.05 (J) Volcano plot of KLF2 versus GFP effects on ATAC (black) and RNA (grey) programs in expanded cells. (K) Accessibility of ATAC programs 8 and 9 in expanded cells across TF perturbations. Points indicate mean ± s.e.m.; GFP (blue) and KLF2 (red) highlighted. (L) UMAP of ATAC program 8 and 9 accessibility (left) with Zheng et al. NME1+ and TCF7+ Tex signatures (right).

Having defined robust regulatory programs, we next assessed how TF overexpression reconfigures them. We quantified the effect of each TF in each condition on program activity, generating a map of 12,616 TF-condition-program connections (Methods, **Extended Table 2**). To compare TFs, we correlated their program-level effects across ATAC programs (**Fig. 2D, lower triangle**) and RNA programs (**Fig. 2D, upper triangle**). As expected, the TFs that clustered separately from wild-type cells (OCT, HOX, and bHLHs; **Extended Data Fig. 4**) produced family-specific effects on chromatin and gene expression (**Fig. 2D**). The OCT factors POU2F2 and POU2F3 induced the strongest changes in program scores (158.84 and 198.68; **Fig. 2D**). An isoform of POU2F2 with an intact DNA binding domain but altered auxiliary domains (P09086-4) showed reduced effects (63.06) but retained highly correlated program-level changes (Pearson R=0.91 with POU2F2; R=0.86 with POU2F3), indicating similar targeting with attenuated activity. Together, these data provide a resource linking TF perturbations to regulatory program activity across CD8^+^ T cell states (**Extended Table 2**).

We sought to identify therapeutically relevant regulatory programs and the TFs that control them. To assess the clinical relevance of the regulatory programs, we collected bulk RNA-seq data from pre-infusion CAR T-cell products, which were subsequently delivered to patients with chronic lymphocytic leukemia^42^ (**Fig. 2E**). Patient samples were scored by expression of RNA programs or genes linked^6^ to ATAC programs. Several programs predicted clinical response (n = 4 associated with positive therapeutic response, n = 11 with negative response; *P* < 0.05) (**Fig. 2F**), with RNA program 1 showing the strongest association with positive response (*P* = 0.001; Fig. 2G). This program was enriched for memory-associated genes (e.g., *SELL, BCL2, IL7R*), elevated in effector-memory cells, and depleted in exhausted states (**Fig. 2C, Extended Data Fig. 5**), suggesting that its activation may oppose exhaustion.

We hypothesized that TFs that upregulate RNA program 1 may be therapeutically relevant targets for immunotherapy. Among all TFs, KLF2 most strongly upregulated RNA Program 1 (**Fig. 2H**). Functional validation confirmed that KLF2 overexpression in T cells enhanced effector capacity in exhausted cells, increasing Granzyme B (40,657 vs 13,424 MFI) and IFN-γ (35.9% vs 8.3% IFN-γ high) production following PMA/ionomycin stimulation relative to GFP controls (**Fig. 2I**). These results suggest KLF2 preserves memory function in exhausted T cells, consistent with knockout phenotypes in mouse models described in prior literature^35^. We next asked whether other regulatory programs could clarify how KLF2 reshapes exhaustion. In expanded cells, KLF2 overexpression preferentially repressed ATAC programs 8 and 9 relative to GFP controls (**Fig. 2J**) and was the strongest repressor of these programs across all TFs (**Fig. 2K**). ATAC programs 8 and 9 correlate with literature signatures corresponding to a precursor exhausted progenitor-like state that sustains terminal exhaustion^43^ (**Fig. 2L**). Together, these findings suggest a mechanism by which KLF2 may limit exhaustion, by restricting entry into the precursor exhausted state.

### TF collaboration enables precise state-specific regulatory control in exhaustion

Having established a functional role for KLF2 in reshaping exhaustion-associated programs, we next sought to define the broader regulatory architecture governing cell state identities in our system. To this end, we applied seq2PRINT^44^, a footprinting-based deep learning framework that infers *de novo* TF binding motifs directly from chromatin accessibility profiles (**Fig. 3A**). We ran seq2PRINT aggregating across 1.63 billion uniquely mapped reads, resulting in the discovery of 289 *de novo* motifs (**Extended Data Fig. 6, Extended Table 3, 4**). Differential accessibility of seq2PRINT *de novo* motifs comparing exhausted and expanded GFP control cells identified established regulators of exhaustion, including motifs corresponding to NR4A^37,45^, NFAT^46^, EGR^47^, and AP-1/BATF^48^ (**Fig. 3B**).

**Figure 3.**
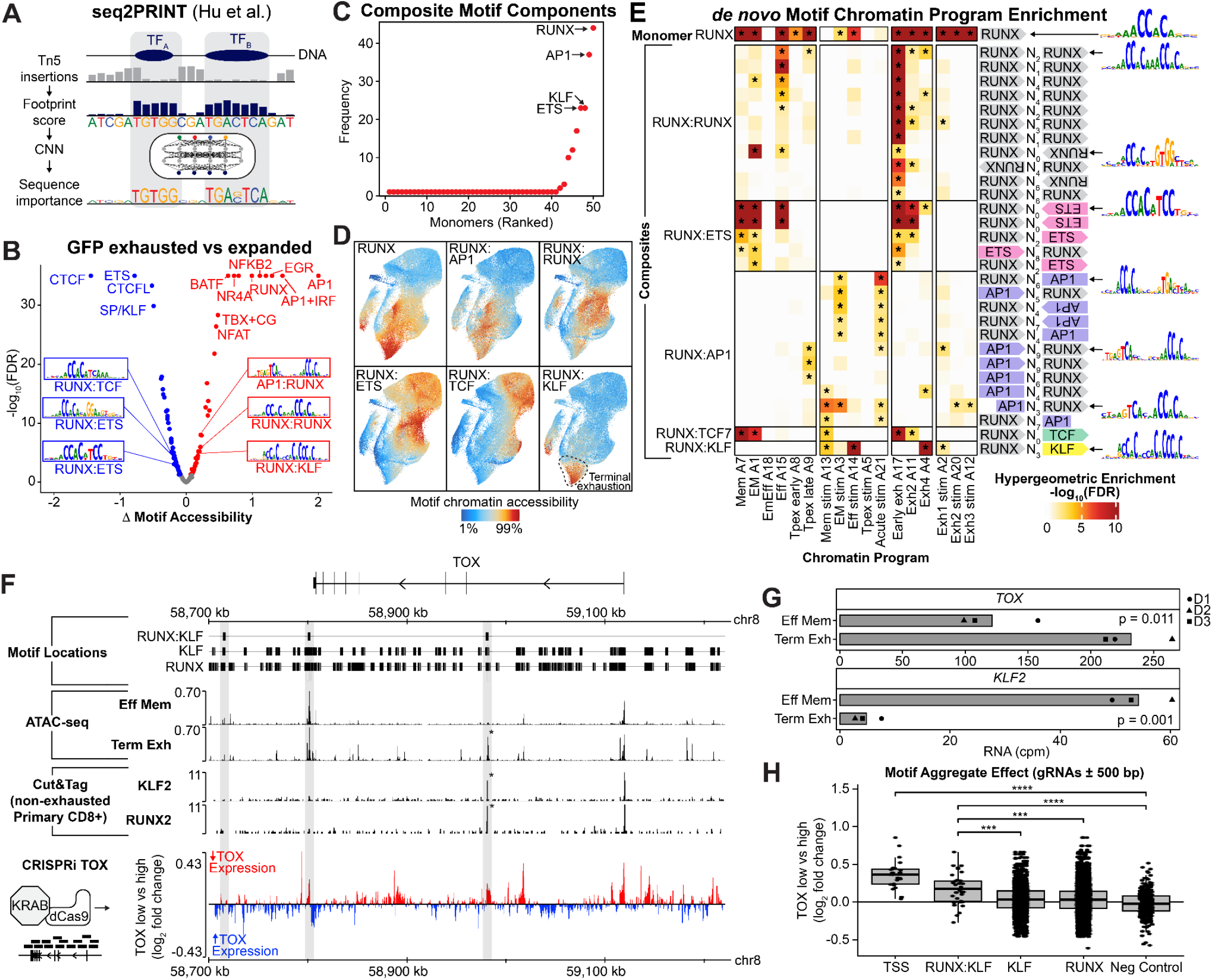
Combinatorial TF motifs define state-specific regulatory programs in exhaustion. (A) Schematic of seq2PRINT^44^, a footprinting-based deep learning framework to identify TF binding motifs. (B) Volcano plot showing differential accessibility of *de novo* motifs in exhausted versus expanded GFP cells. (C) Dot plot showing the frequency of each monomer motif occurrence in composite motifs, ranked from lowest to highest frequency. (D) UMAP of unstimulated cells colored by chromatin accessibility of RUNX monomers, homodimers, and heterodimers. (E) Heatmap of RUNX monomer and composite *de novo* motif (rows) enrichment within chromatin programs (columns), colored by -log_10_ FDR of hypergeometric enrichment. Motif spacing and orientations are indicated on the right, with example motif PWMs highlighted. * indicates FDR < 0.05. (F) Genomic track of the TOX locus highlighting epigenetic features. Motif Locations: Genomic positions of *de novo* motifs corresponding to RUNX:KLF composite sites, as well as RUNX and KLF monomer motifs within the TOX locus. RUNX:KLF motifs are highlighted in grey. ATAC-seq: Pseudobulked chromatin accessibility tracks from effector memory (Eff Mem) and terminal exhaustion (Term Exh) cells across the TOX locus, normalized to insertions per million. Cut&Tag: Profiles of HA-tagged KLF2 and RUNX2 localization within the TOX locus in healthy primary human CD8^+^ T cells, normalized to fragments per million. CRISPRi TOX: (left) Schematic of CRISPRi guide tiling across regulatory elements at the TOX locus. (right) Genomic track of the log_2_ fold change of CRISPRi guides targeting the TOX locus. Red indicates regions that decrease TOX expression; blue indicates regions that increase TOX expression. (G) RNA expression counts per million (cpm) of *TOX* and *KLF2*, pseudobulked by donor for effector memory (Eff Mem) and terminally exhausted (Term Exh) cells. (H) Mean log_2_ fold change in TOX expression for guides targeting 1-kb windows centered on the TSS, *de novo* motif sites, and negative controls.

We identified numerous composite *de novo* motifs (n=131), defined by adjacent or overlapping binding sites for multiple TFs (**Extended Table 3, 4**). Compared to monomeric motifs (median 29,153 binding sites per motif, n=53 monomeric motifs), composite motifs were associated with fewer binding sites overall (median 1,575 per motif), yet remained significantly enriched across all chromatin programs (**Extended Data Fig. 6**). Prior work has largely described TF cooperation as the interaction of unique pairs of factors^49^. Surprisingly, we found that composite motifs were largely skewed in their composition (**Fig. 3C**). To define the composition of these motifs, we analyzed their constituent monomeric TF pairs (n=125) and trios (n=6). TF collaboration was highly concentrated among 4 factors (RUNX, AP1, KLF, and ETS), comprising 68% of all composite motifs. Specifically, the RUNX motif appeared 44 times within the composite motifs, followed by AP1 (n=24), KLF (n=23), and ETS (n=23) (**Fig. 3C**). This outstanding degree of motif co-occurrence led us to conclude that RUNX is a master collaborator, likely enabling cells to specify distinct gene expression programs with a limited set of factors.

Interestingly, RUNX-containing composite motifs exhibited bidirectional changes in accessibility in exhaustion (**Fig. 3B**). Although RUNX proteins are classically described as obligate heterodimers with the co-activator CBFβ, accumulating evidence indicates that RUNX family members homodimerize^50,51^ and heterodimerize^52–54^ with other TFs. The diverse RUNX composite motifs were highly specific to distinct cellular states (**Fig. 3D**). Furthermore, we found that RUNX composite motifs were specifically enriched in distinct chromatin programs (**Fig. 3E, Extended Data Fig. 6**). More broadly, composite motifs of any TF had greater state specificity than monomers (Mean Gini coefficient 0.37 monomer, 0.50 composite, *P* = 1x10^-4^, **Extended Data Fig. 6**). These findings support a model^49^ in which combinatorial motifs enable cells to precisely activate cellular states using a limited set of TFs.

To further explore RUNX as a “master collaborator”, we asked whether composite motif orientations influence binding partners. Using Cistrome^55^, we quantified TF ChIP-seq co-occupancy across distinct orientations and spacings of RUNX homo- and heterodimer motifs (**Extended Data Fig. 6)**. Interestingly, RUNX homodimer configurations exhibited orientation- and spacing-dependent differences in CBFβ co-occupancy. Heterodimer partners likewise showed distinct co-binding patterns: RUNX:ETS motifs were strongly enriched for CBFβ occupancy (GIGGLE score 1166.182), whereas RUNX:KLF motifs showed no significant overlap (**Extended Data Fig. 6**). Together, these findings suggest that composite motif orientation and spacing constrain co-factor recruitment.

Returning to exhaustion, we found that accessibility of the RUNX:KLF composite motif was specifically accessible in terminally exhausted cells (**Fig. 3D**) and enriched in the chromatin program underpinning terminal exhaustion (**Fig. 3E**). Consistent with the literature, KLF2 expression decreased in exhaustion (*P* = 0.001, **Fig. 3G**). KLF2 acts as a repressor in T cells^35^, leading us to hypothesize KLF2 may repress exhaustion programs. Although the RUNX:KLF composite motif is rare genome-wide, present in only 0.86% of regulatory regions (2,970 of 345,953 peaks), it occurs three times within the TOX locus (**Fig. 3F**). Chromatin accessibility of one of these regions significantly increased over exhaustion (**Fig. 3F)**. CUT&Tag profiling with overexpressed HA-tagged KLF2 and RUNX2 confirmed that the same locus was bound by both factors in primary human CD8^+^ T cells, supporting cooperative binding at the composite motif (**Fig. 3F, Extended Data Fig. 7)**. We chose RUNX2 due to its increased expression in exhaustion and its enrichment in terminal exhaustion programs (**Extended Data Fig. 5**). This motivated us to return to the original screen and analyze the RUNX TF family and their isoforms. All isoforms of RUNX2 decreased TOX expression in the exhausted screen (n=4 isoforms, **Extended Table 1**), whereas RUNX1 and RUNX3 had bidirectional effects. Together, these data support a model in which RUNX2:KLF2 co-binding modulates chromatin accessibility at exhaustion-associated regulatory elements to control TOX expression.

Because chromatin accessibility and TF colocalization are only correlative with gene expression, we sought to directly test whether the identified regulatory elements causally influence TOX expression. We performed a CRISPR interference (CRISPRi) tiling screen across the TOX locus (n=6,733 targeting guides, n=305 controls, **Extended Data Fig. 7**). Guide RNAs were introduced into primary CD8^+^ T cells, followed by chronic stimulation to induce exhaustion (**Fig. 3F**). Cells were subsequently sorted into TOX^HIGH^ and TOX^LOW^ populations based on protein abundance, and guide enrichment between bins was quantified to assess the regulatory contribution of each targeted region (**Extended Data Fig. 7)**. Guides targeting two of the RUNX:KLF containing motifs significantly reduced TOX expression (log_2_ fold change = 0.66, *P* = 0.0003; log_2_ fold change = 0.46, *P* = 0.005) (**Extended Data Fig. 7**). Aggregating guides within 1-kb windows centered on individual motifs revealed that targeting RUNX or KLF monomeric sites produced modest enrichment in the TOX^LOW^ population (9.3% and 9.8%, respectively, relative to TSS-targeting guides), whereas targeting the RUNX:KLF composite motif yielded substantially stronger effects (47% of TSS magnitude; **Fig. 3H**). Among all motifs in the TOX locus (n=114 with >5 guides), only NRF exceeded the RUNX:KLF effect (66.4% of TSS magnitude), consistent with recent work implicating NRF2 in T cell exhaustion^56^ (**Extended Data Fig. 7**). Taken together, these results indicate that RUNX:KLF composite elements more precisely control TOX protein levels than monomeric binding sites alone.

Here we identify RUNX as a hub of collaborative TF activity in CD8^+^ T cell states. We identify RUNX2:KLF2 collaboration as a defining feature of exhaustion, and find that the motif exerts specific control over TOX expression. Together, our data suggest that KLF2 interacts with RUNX2 to control exhaustion-associated programs.

### RUNX2:KLF2 interaction modulates exhaustion-associated gene expression

Together, these results establish RUNX2:KLF2 collaboration as a central regulator of TOX expression, prompting us to investigate the molecular basis of this interaction. We performed an *in vitro* footprinting assay to measure the DNA binding and cooperativity of RUNX2 and KLF2 (**Fig. 4A**). Briefly, full-length RUNX2 and KLF2 proteins were incubated with 152 kb of human DNA, and Tn5 was used to footprint DNA and measure TF occupancy^44,57^. KLF2 bound the KLF2 target motif (n=128/164 KLF2 motifs bound with footprint *P* < 0.05, **Fig. 4B**). In contrast, RUNX2 alone exhibited limited binding (n=1/13 bound to monomeric sites, n=3/11 bound to dimer sites), consistent with known limitations of RUNX binding in the absence of co-factors such as CBFB^58^ (**Fig. 4B**). When RUNX2 and KLF2 were combined, TF occupancy at monomeric sites remained largely unchanged, however composite RUNX:KLF motifs exhibited significantly increased binding (n=16/25 RUNX+KLF score increased more than 5% versus KLF alone) (**Fig. 4B, C**). Interestingly, this enhancement was dependent on motif spacing: no cooperativity was observed when the two motifs were directly adjacent (n=0/4) (**Fig. 4B**). Together, this demonstrates that RUNX2 and KLF2 collaborate in a DNA sequence-dependent manner.

**Figure 4.**
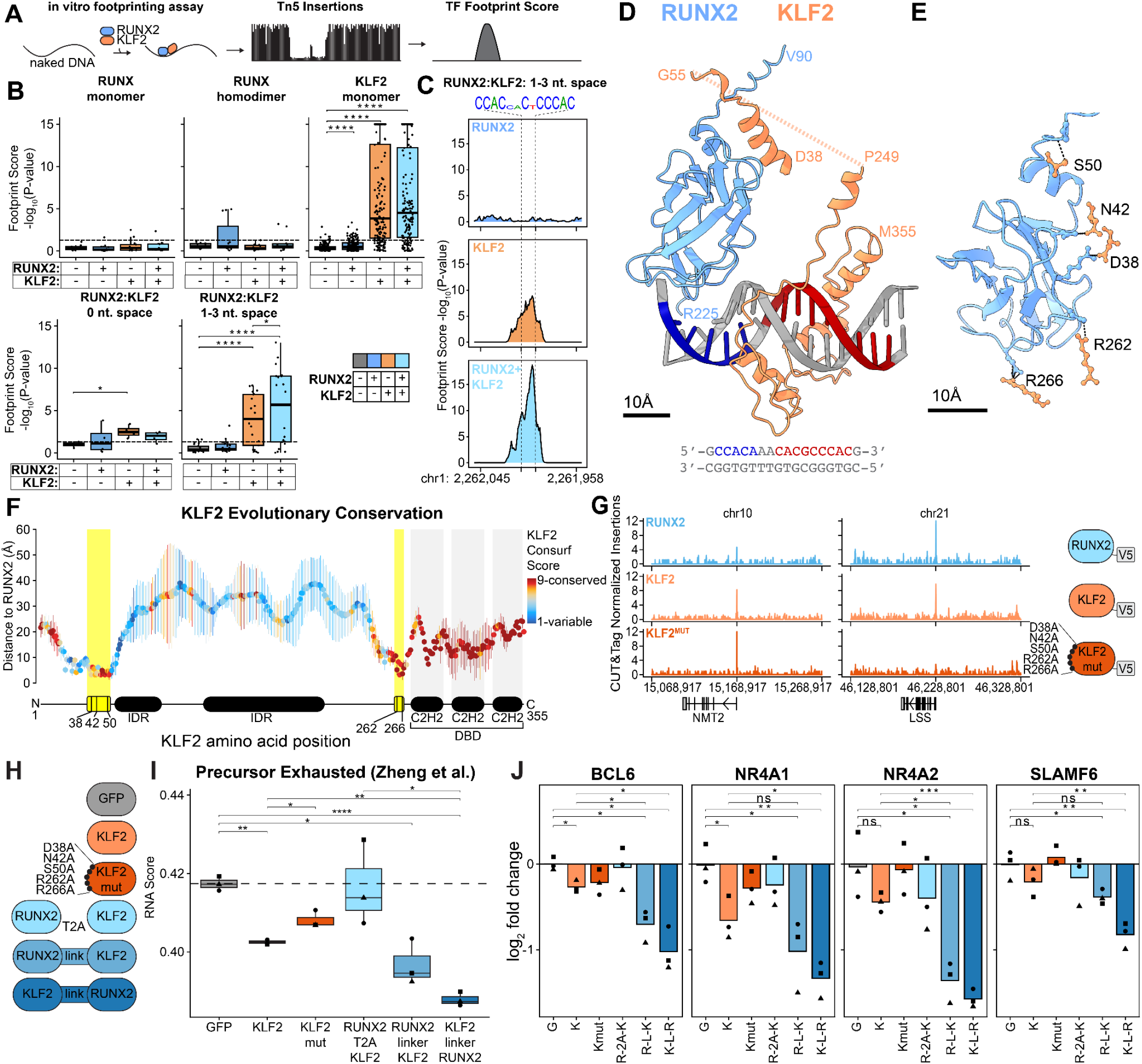
RUNX2:KLF2 interaction at composite motifs modulates exhaustion-associated gene expression. (A) Schematic of the *in vitro* footprinting assay. Naked DNA is incubated with purified TFs and transposed with Tn5 to assess TF binding-dependent protection. (B) Footprint scores at RUNX and KLF monomeric, homodimeric, and heterodimeric motifs (RUNX:KLF is CCACN_x_CNCCCAC, with x = nucleotide spacing), measured as -log10 *P* of TF binding, across: no TF added (grey), RUNX2 (blue), KLF2 (orange), and RUNX2+KLF2 (light blue) conditions. *P* values from paired two-sided T-test. (C) Representative footprint profiles at a composite motif for RUNX2, KLF2, and combined RUNX2+KLF2 conditions. (D) AlphaFold3 (AF3) model of RUNX2 (blue) and KLF2 (orange) bound to the composite RUNX:KLF *de novo* DNA motif, with RUNX2 and KLF2 individual motifs highlighted in blue and red, respectively. Scale bar indicates 10 Å. (E) AF3-predicted interface between RUNX2 and KLF2, with KLF2 residues labeled. Dashed lines indicate hydrogen bonds (N42, S50, R262) and polar contacts (D38, R266). Scale bar indicates 10 Å. (F) Heavy-atom distances between KLF2 residues and their nearest RUNX2 residue across five AF3 predictions (mean ± s.d.). Residues are colored by evolutionary conservation (ConSurf). Yellow shading marks residues forming hydrogen bonds or polar contacts; black boxes denote the intrinsically disordered domains (IDR) and the DNA binding domain (DBD), including three C2H2 zinc fingers (C2H2 I–III). (G) CUT&Tag genomic tracks of RUNX2, KLF2, and KLF2^MUT^ occupancy at NMT2 (left) and LSS (right). Insertions are normalized to fragments per million. (H) Schematic of constructs used for overexpression experiments, including GFP control, KLF2, KLF2^MUT^, and RUNX2+KLF2, and tethered variants. (I) Precursor exhaustion RNA score (Zheng et al.) across overexpression conditions in (H), indicating modulation of exhaustion-associated transcriptional programs. *P* values from two-sided T-test *: P ≤ 0.05, **: P ≤ 0.01, ***: P ≤ 0.001. (J) Log_2_ fold change in individual gene expression relative to the GFP mean across overexpression conditions: G (GFP), K (KLF2), Kmut (KLF^MUT^), R-2A-K (RUNX2-T2A-KLF2), K-2A-R (KLF2-T2A-RUNX2). *P* values from two-sided T-test: n.s.: P > 0.05, *: P ≤ 0.05, **: P ≤ 0.01, ***: P ≤ 0.001.

To understand the structural basis of this cooperativity, we used AlphaFold 3 (AF3)^59^ to predict the structure of RUNX2:KLF2 bound at the composite DNA motif (**Fig. 4D**). The structure suggested a direct interaction between RUNX2 and KLF2 and identified specific hydrogen bonds (KLF2 positions: N42, S50, R262) and polar contacts (D38, R266) mediating the protein–protein interface (**Fig. 4E**). Across five independent AF3 iterations, these residues consistently remained in close proximity with minimal structural variance (**Fig. 4F**). Interface residues displayed elevated evolutionary conservation in both KLF2 (**Fig. 4F**) and RUNX2 (**Extended Data Fig. 9**), supporting their functional significance.

To assess the functional relevance of this interface, we generated a KLF2 mutant in which all of the predicted interacting residues (N42, S50, R262, D38, R266) were substituted with alanines (KLF2^MUT^). CUT&Tag profiling of HA-tagged KLF2^MUT^ in primary human CD8^+^ T cells demonstrated that global DNA binding was largely preserved relative to wild-type KLF2 (KLF2^WT^) (**Fig. 4G**), and binding sites remained enriched for canonical KLF motifs (**Extended Data Fig. 8**). However, KLF2^MUT^-bound regions exhibited depletion of overlapping binding with RUNX2 motifs (n=82 of 234 shared binding sites lost), consistent with impaired cooperative occupancy (**Fig. 4G, Extended Data Fig. 8**). Interestingly, KLF2^MUT^ was the most significantly depleted relative to KLF2^WT^ at the promoter of LSS (Lanosterol synthase), a key enzyme in the mevalonate-cholesterol pathway that is implicated in CD8^+^ tissue residency^60^, raising the possibility that RUNX2:KLF2 cooperativity may contribute to this transcriptional program (**Fig. 4G**). Overall, these results indicate that the predicted RUNX2:KLF2 interface is dispensable for DNA binding but required for certain instances of RUNX-associated chromatin engagement.

Given that KLF2 overexpression decreased precursor exhaustion-associated programs (**Fig. 2J**), we next asked whether an interaction with RUNX2 was required for this effect. To test this, we overexpressed KLF2^WT^ and KLF2^MUT^ in primary human CD8^+^ T cells and scored cells using a literature-derived precursor exhaustion RNA signature^41^. As expected, overexpression of KLF2^WT^ in primary human CD8^+^ T cells led to a significantly reduced precursor exhaustion RNA score (*P* = 0.0025). Mutating predicted residues of RUNX2 interaction (KLF2^MUT^) attenuated this phenotype (*P* = 0.039 vs KLF2^WT^) (**Fig. 4I**). Thus, disruption of the predicted RUNX2:KLF2 interface abrogates the ability of KLF2 to repress exhaustion-associated transcriptional programs.

Finally, we asked whether enforced RUNX2:KLF2 interaction is sufficient to amplify this phenotype. The simultaneous binding of two TFs at a composite motif is extremely rare within the genome and is dictated by the TF concentration, genomic search space, kinetics, and thermodynamic binding affinities^3^. We hypothesized we could synthetically increase composite motif binding by tethering RUNX2 and KLF2 together using a flexible amino acid linker, thereby reducing the genomic search space and synthetically increasing effective local concentration. The tethered RUNX2-KLF2 constructs produced an even more pronounced reduction in precursor exhaustion RNA signatures compared to wild-type KLF2 alone (*P* = 0.018, *P* = 0.00004) and to a RUNX2-T2A-KLF2 construct, which produces two separate proteins (*P* = 0.042) (**Fig. 4I**). This effect extended beyond aggregate scores, as individual exhaustion-associated genes, including BCL6, NR4A1, NR4A2, and SLAMF6, exhibited concordant significant decreases (**Fig. 4J**).

Collectively, integrative structural, genomic, and functional analyses uncover a novel RUNX2:KLF2 collaboration at composite motifs that targets exhaustion-associated gene expression programs. Leveraging our combined experimental and computational framework, we show that enforced RUNX2:KLF2 tethering is sufficient to amplify this program. These findings define TF collaboration as a programmable mechanism for controlling cell state, establishing a new paradigm for engineering cellular identity.

## Discussion

Understanding how TFs specify immune cell identity has remained a central question in immunology. Although numerous regulators of CD8^+^ T cell activation, memory, and exhaustion have been identified, the organizing principles by which these factors collectively encode cell state have been less clear. Here, by integrating genome-wide perturbation with multimodal single-cell profiling, functional regulatory element dissection, and structural modeling, we identify RUNX as a master collaborator that governs CD8^+^ T cell states. More broadly, this reveals a category of TFs with pervasive partners that might serve as core hubs modulating cell state identity within a given cell type.

It is well appreciated that TFs cooperate and that co-binding confers regulatory specificity^61^. Classical reprogramming studies demonstrated that defined TF cocktails can redirect cell fate^14–16^, and more recent epigenomic analyses have revealed a broad repertoire of structured TF–TF interactions embedded within chromatin landscapes^62–65^. Our study advances this framework from observation to engineering. We show that the cooperative interactions that define endogenous regulatory systems can be systematically discovered and rationally engineered. Mutating the protein-protein interface establishes that a defined RUNX2:KLF2 protein-protein surface is required for KLF2’s functional repression of exhaustion programs, and synthetic tethering demonstrates that enforced proximity is sufficient to amplify this regulatory effect.

Collaborative TF binding distinctly partitions CD8^+^ T cell states. By physically linking RUNX2 and KLF2 into a single functional unit, we convert this composite logic into a higher efficiency regulatory actuator: the tethered construct more strongly suppresses precursor exhaustion programs and more robustly reinstates a therapeutically favorable state associated with improved CAR-T response than KLF2 alone. This provides a mechanistic rationale for extending prior work on KLF2 as a regulator of T cell function^35^ and positions RUNX2:KLF2 as a more potent and precise engineering candidate. More broadly, these findings outline a generalizable paradigm for cell engineering in which composite motif architectures are targeted through synthetic TF tethering, enabling rational access to gene programs that are otherwise refractory to modulation by single factors.

## Acknowledgements

We are grateful to the Buenrostro and Gersbach laboratories for helpful feedback throughout this project. We would like to thank Ena Oreskovic for her expertise in immunology, and Zhi-Jie Cao for computational help. We greatly appreciate Surya Nagaraja, Sidu Jena, Zack Chiang, and Henry Bushnell for helpful discussions. We would like to thank Tomer Rotstein and Rachel Conover for help on validation experiments. We would like to thank the IGVF Immune Focus group for feedback and support. We would like to thank Siddarth Wekhande, Nina Farrell, Fabiana Duarte, Eugenio Mattei, Liz Gaskell, Katie Irish, Noam Shoresh, Amelia Hall, Chuck Epstein, Bradley Bernstein, Alejandro Barrera, and Ruhi Rai for support related to the NIH IGVF Consortium.

## Author contributions

C.M.A. and A.G. performed the genome-wide MORF screen, prepared cells for single-cell screens, performed flow-based KLF2 validation, and performed the CRISPRi screen. C.T.H. developed the *in vitro* footprinting and analysis. R.Z. performed seq2PRINT and cisTopic modeling. M.A.H. contributed to project conception and analysis. J.K. and R.E.S. performed CD8^+^ T cell isolation and electroporation. R.E.S. performed all other experiments and analysis. C.A.G. and J.D.B. supervised all aspects of the study. R.E.S. and J.D.B. wrote the manuscript with input from all authors.

## Data and materials availability

As of 4/19/2026, raw and processed data have been provided to the Impact of Genomic Variation on Function (IGVF) consortium and may be accessed at https://data.igvf.org/analysis-sets/IGVFDS7404QFES/. CRISPRi data are also provided to IGVF at https://data.igvf.org/analysis-sets/IGVFDS4798PRUI/. Additional data for bulk assays can be found under the superseries GSE325697, with the following: bulk RNA-seq, CUT&Tag, in vitro footprinting, TF overexpression screen. This preprint will soon be updated accordingly. Access to data will be made available upon request. All code is being made available on GitHub: https://github.com/rachel-e-savage/savage-perturbshare

## Methods

### Isolation and culture of primary human T cells

Human CD8^+^ T cells were isolated from three distinct donors (StemCell Tech, cat no. 70500) using negative selection human CD8 isolation kits (StemCell Tech, cat no. 17953). T cells were activated using anti-CD3/CD28 Dynabeads (Gibco, cat no. 11132D). For all experiments, T cells were cultured in PRIME-XV T cell Expansion XSFM (FujiFilm, 91141-1L) supplemented with 5% human platelet lysate (Compass Biomed, PLSA500), 100 U ml^−1^ penicillin, 100 μg ml^−1^ streptomycin (Gibco, cat no. 15140122), and 100 U ml^-1^ of human IL-2 (Peprotech, 200-02-1MG). T cells were activated with a 3:1 ratio of CD3/CD28 dynabeads to T cells and maintained at 1–2 × 106 cells ml−1 unless otherwise indicated. For chronic stimulation experiments, dynabeads were removed using a magnet, and fresh dynabeads at a 3:1 Dynabead:T cell ratio were added.

### Lentivirus generation and transduction of primary human T cells

For all lentivirus generation, a transfection protocol from Schmidt et al. was used to ensure high lentiviral titer^19^. In brief, HEK-293Ts were transfected with lentiviral packaging, envelope, and vector of interest plasmids. Lentiviral supernatants were harvested 24 hours and 48 hours after transfection, with a media refresh between harvests. Lentiviral supernatant was centrifuged at 600g for 10 min to remove cellular debris and concentrated to 50–100× the initial concentration using Lenti-X Concentrator (Takara Bio, cat no. 631232). T cells were transduced at 5–10% v/v of concentrated lentivirus at 24 h post-activation. For co-transduction experiments, T cells were transduced at 5-10% v/v of concentrated lentivirus 24 h post activation.

### mORF TF-wide screen

Day 0: CD8^+^ T cells were transduced (n=3 biological replicates) with the mORF pooled lentiviral library (Addgene #192821) at 10% v/v, 24 hours after isolation and stimulation with anti-CD3-CD28 Dynabeads (see Isolation and culture of primary human T cells), and expanded for 3 days in standard media. Day 3: Cells were selected with puromycin at 1 µg/mL for two days. Day 5: Dynabeads were removed, and cells were split into two conditions: expanded and exhausted. Both conditions were supplemented with 0.5 ug/mL of puromycin, and the exhaustion condition was additionally restimulated with fresh dynabeads at a 3:1 concentration. Day 7: both conditions were replenished with standard media with no puromycin. Dynabeads were removed and replenished for the exhaustion condition at a 3:1 concentration. Day 9: Expanded cells were fix/permed, and Day 10: exhausted cells were fixed/permed following manufacturer protocol (FOXP3 TF staining kit, Thermo, cat no. 00-5523-00), and stained for TOX (Miltenyi Biotec, cat no. 130-118-335) at 1:100 dilution. Transduced cells were sorted for top and bottom 10% TOX expressing cells using an SH800 FACS Cell Sorter (Sony Biotechnology) for subsequent genomic DNA isolation for mORF barcode library construction and sequencing. All replicates were maintained and sorted at a minimum of 100x coverage. Cells were maintained at 1–2 × 10^6^ cells ml^−1^ for all expansion steps unless otherwise indicated.

### Genomic DNA isolation, mORF barcode PCR and sequencing mORF libraries

Genomic DNA was isolated using the PicoPure DNA extraction kit (Applied Biosystems, cat no. KIT0103) following manufacturer recommendations and incubating the samples for 65 C overnight for reverse crosslinking. Genomic DNA was split across 100 μl PCR reactions (22 cycles at 98 °C for 10 s, 60 °C for 10 s, and 72 °C for 25 s) with NEBNEXT 2× Master Mix (NEB, cat no. M0541L) and up to 1 μg of genomic DNA per reaction. PCRs were pooled together for each sample and purified using double-sided magnetic Ampure XP bead selection (Beckman Coulter, cat no. A63882) at 0.5× and 1.0×. Libraries were run on a High Sensitivity D1000 tape (Agilent, cat no. 5067-5584) to confirm amplicon size and quantified using Qubit’s dsDNA High Sensitivity assay (Invitrogen, cat no. Q32851). Libraries were diluted to 2 nM, pooled together at equal volumes, and sequenced using MiSeq Reagent Kit v2 50 cycles (Illumina, cat no. MS-102-2001). Primers are listed in Supplemental Oligo Table under TF_mORF_NGS_FW, TF_mORF_NGS_BC_REV.

### Processing mORF sequencing and mORF barcode analysis for FACS-based screens

FASTQ files were aligned to custom indexes for each mORF library (generated from the bowtie2-build function) using Bowtie 2. Counts for each mORF barcode were extracted and individual TF enrichment was determined using DESeq2 to compare mORF abundance between groups for each screen. Full DESeq2 results for mORF screens are presented in Extended Table 1.

### minimORF subpooled perturb-SHARE-seq screen

Plasmids of all mORF TFs that were significantly enriched and depleted from the exhaustion TF-wide screen and were not significant in the expanded stimulation screen were pooled equimolarly. Individual plasmids were purchased from Addgene (see Extended Table 1 for the MORF ids included in perturb-SHARE-seq). The subpool of “exhaustion-unique” TFs was then sequenced and analyzed as the FACS-based screen (see “Genomic DNA isolation, mORF barcode PCR and sequencing mORF libraries” and “Processing mORF sequencing and mORF barcode analysis for FACS-based screens”).

In brief, the lentiviral screen followed the same timeline as the mORF TF-wide screen until day 9. At day 9, the expanded conditions were expanded for an additional day alongside the exhausted condition. On day 10, expanded and exhausted cells were split in half, and half were stimulated with 1x T cell stimulation cocktail (Invitrogen, cat no. 00-4970-03) for two hours and fixed for further processing (see “Fixation” and “perturb-SHARE-Seq”). The other half were mock stimulated with an equal volume of ethanol for two hours, and fixed for further processing. Protocol published on protocols.io at the following DOI: https://dx.doi.org/10.17504/protocols.io.6qpvrbwjplmk/v1

### CRISPRi tiling screens

CD8^+^ T cells were transduced 24 hours after activation with all-in-one lentiviral vector encoding for dSpCas9–KRAB–2A–Thy1.1 and gRNAs targeting the TOX locus ±100 kb (n = 3 biological replicates). Cells were restimulated with Dynabeads every two days after transduction for a total of three restimulations, and were expanded for a final four days. Cells were then stained using a Thy1.1:PE antibody (Biolegend, cat no. 109005) at a 1:300 dilution, fix/permed following manufacturer protocol (FOXP3 TF staining kit, Thermo, cat no. 00-5523-00), and stained for TOX (Miltenyi Biotec, cat no. 130-118-335) at 1:100 dilution. Transduced cells were gated based on Thy1.1 signal and top and bottom 10% TOX expressing cells were sorted for subsequent genomic DNA isolation for gRNA library construction and sequencing. All replicates were maintained and sorted at a minimum of 100× coverage. Cells were maintained at 1–2 × 10^6^ cells ml^−1^ for all expansion steps unless otherwise indicated. Protocol also published on protocols.io at the following DOI: https://dx.doi.org/10.17504/protocols.io.rm7vz4e72lx1/v1

### CRISPRi genomic DNA isolation, gDNA PCR and sequencing gRNA libraries

Genomic DNA was isolated using the PicoPure DNA extraction kit (Applied Biosystems, cat no. KIT0103) following manufacturer recommendations and incubating the samples for 65 C overnight for reverse crosslinking. Genomic DNA was split across 100 μl PCR reactions (25 cycles at 98 °C for 10 s, 60 °C for 30 s, and 72 °C for 20 s) with Q5 2× Master Mix (NEB, cat no. M0492L) and up to 1 μg of genomic DNA per reaction. PCRs were pooled together for each sample and purified using double-sided magnetic Ampure XP bead selection (Beckman Coulter, cat no. A63882) at 0.6× and 1.8×. Libraries were run on a High Sensitivity D1000 tape (Agilent, cat no. 5067-5584) to confirm amplicon size and quantified using Qubit’s dsDNA High Sensitivity assay (Invitrogen, cat no. Q32851). Libraries were diluted to 2 nM, pooled together at equal volumes, and sequenced using MiSeq Reagent Kit v2 50 cycles (Illumina, cat no. MS-102-2001). Primers are listed in Supplemental Oligo Table under CRISPRi_NGS_i5_amp, CRISPRi_NGS_BC_NGS_i7_rev.

### PMA/IO functional T cell stimulation assay

CD8^+^ T cells were transduced (n = 3 biological replicates, same donors as the screens) with KLF2 mORF lentivirus (Addgene #142715) at 10% v/v, 24 hours after isolation and stimulation with anti-CD3-CD8 Dynabeads (see Isolation and culture of primary human T cells), and expanded for 3 days in standard media. Day 3: Cells were selected with puromycin at 1 µg/mL for two days. Day 5: Dynabeads were removed, and cells were split into two conditions: expanded and exhausted. Both conditions were supplemented with 0.5 ug/mL of puromycin, and the exhausted condition was additionally restimulated with fresh dynabeads at a 3:1 concentration. Day 7: both conditions were replenished with standard media with no puromycin. Dynabeads were removed and replenished for the exhausted condition at a 3:1 concentration. Day 10: Expanded and exhausted cells were stimulated with 1x T cell stimulation cocktail (Invitrogen, cat no. 00-4970-03) for two hours, stained for CD3 (Invitrogen, cat no. 69-0038-41), CD8 (Invitrogen, cat no. H003T02B06-A), CD4 (Invitrogen, cat no. H001T02B09-A), LAG3 (Invitrogen, cat no. 406-2239-41), TIGIT (Invitrogen, cat no. 12-9500-41), CD69 (Invitrogen, cat no. 47-0699-42), 4-1BB (Invitrogen, cat no. 62-1379-42), PD-1 (cat no. 407-2799-41) (see Flow cytometry and surface marker staining for protocol), fix/permed following manufacturer protocol (Intracellular Fixation & Permeabilization Buffer Set, Invitrogen, cat no. 88-8824-00), and stained for IFN-γ (Biolegend, cat no. 506518) and Granzyme B (Invitrogen, cat no. GRB05). Cells were maintained at 1–2 × 10^6^ cells ml^−1^ for all expansion steps unless otherwise indicated.

### Flow cytometry and surface marker staining

For analyzing and sorting the ORF TF-ome FACS based screens, an SH800 FACS Cell Sorter (Sony Biotechnology) was used. For the PMA/IO stimulation assay, an NXT Attune flow cytometer (Thermo) was used for analysis of expression of IFN-γ and Granzyme B. To stain surface markers, cells were collected, spun down at 300*g* for 5 min, resuspended in Cell Staining Buffer (Biolegend, cat no. 420201) with the manufacturer recommended antibody dilutions and incubated for 30 min at 4 °C on a rocker. Cells were then washed with flow cytometry buffer, spun down at 300*g* for 5 min and resuspended in flow buffer for cell sorting or analysis. Fluorescent minus one (FMO) controls were used to set appropriate gates for all flow panels. For antibody staining of all intracellular markers the Intracellular Fixation & Permeabilization buffer (Invitrogen, cat no. 88-8824-00) set was used following manufacturer’s protocol. In brief, after surface staining, cells are spun down, resuspended in fixation buffer, and incubated at room temperature for 30 minutes. Afterwards cells were washed with permeabilization buffer, resuspended in permeabilization buffer with manufacturer-recommended antibody dilutions, and stained for 30 minutes at room temperature. Cells were washed twice with permeabilization buffer, resuspended in flow buffer and analyzed via flow cytometry. All relevant gating/analysis was conducted using Flow Jo V10.10.0.

### Fixation

Fixation was performed as described by Hu et al.^44^ in a final concentration of 0.2% formaldehyde. Following fixation quenching and washing, cell pellets were resuspended in BamBanker (BB05) and frozen in a Mr. Frosty™ Freezing Container to -80C.

### perturb-SHARE–seq

Frozen fixed cell pellets were thawed by an equal volumetric addition of 4C PBS-RI. SHARE-seq with additional RNA capture (through poly-adenylation of RNA transcripts prior to reverse transcription) was performed as described in Hu et al. using preassembled Tn5 (seqWell, Tagify SHARE-seq Reagent). The full protocol is published at https://dx.doi.org/10.17504/protocols.io.81wgbx1oylpk/v5. The following modifications were made in order to capture the TF-barcoded transcript. During reverse transcription, 1 uL of 100 uM MORF-RT-Primer was added in addition to the standard 5 uL of 100 uM RT primer. Following template switching, during RNA PCR, 0.2uL of 25uM MORF-RNA-Primer (combining equimolar ratios of 8 primers with variable spacing) was spiked in. Following PCR amplification, products were purified using 0.8x Ampure. The purified product was split into two. One half was processed as the sc-RNA-seq library, as described by Hu et al. The other half of the purified product was additionally amplified using 0.8 uL of 25uM P7 primer and 0.8uL of 25uM MORF-RNA-Primer. This PCR product was purified and amplified a final time with P7 and a barcoded TruSeq primer, treated with ExoI, and Ampure purified (0.8X). Primer sequences are located in Supplemental Oligo Table.

### Quantification and sequencing

scATAC-seq, scRNA-seq, and scTF-barcode-seq libraries were quantified with the KAPA Library Quantification Kit and pooled for sequencing. Single-cell libraries were sequenced on the Nova-seq platform (Illumina) with the following parameters: Read 1: 100; Read 2: 100; Index 1 (i7): 99; Index 2 (i5): 8.

### Donor calling

Frozen fixed cell pellets from each donor were thawed by an equal volumetric addition of 4C PBS-RI. Nuclei were isolated and bulk ATAC-seq was performed as described in Buenrostro et al. using preassembled Tn5 (seqWell, Tagify SHARE-seq Reagent). Donor genotypes were called with GATK v4.1.2.0 HaplotypeCaller and jointly genotyped; single cells were assigned to donors using Drop-seq AssignCellsToSamples.

### Plasmid libraries

MORF library was purchased from Addgene (137000). For validation experiments, TFs were individually synthesized, inserted, and sequence verified by TWIST onto a pLEX_307 (Addgene #41392) backbone. Plasmid structure is depicted in Extended Data Figure 9C.

### CD8^+^ electroporation

Human CD8^+^ T cells were isolated using EasySep™ Human CD8^+^ T Cell Isolation Kit (Stem Cell Technologies #17953) from 100M human PBMCs (Stem Cell Technology #70025). Cells were plated at 1M/mL in ImmunoCult™-XF T Cell Expansion Medium (Stem Cell Technology #10981) with IL-2 and expanded for 4 days. Cells were electroporated using the P3 Primary Cell 4D-Nucleofector® X Kit S (Lonza #V4XP-3032) following the manufacturer’s recommended protocol for human T cells. Cells were replated to 1M/mL and cultured for an additional 48 hours prior to CUT&Tag and RNA-seq experiments.

### CUT&Tag

CUT&Tag was performed following EpiCypher Version 2.0 buffers, and Version 2.1 protocol. The following antibodies were used: CUTANA™ pAG-Tn5 (15-1017) and V5 antibodies (Cell Signaling Technology Inc D3H8Q), HA antibody (EpiCypher 13-2010). Following pAG-Tn5, samples were processed as in Buenrostro et al.

### Bulk RNA-seq

100k cells were resuspended in Zymo DNA/RNA Shield (Zymo R1100) and processed via Plasmidsaurus RNA-seq, using 3’ Poly-A sequencing, capturing ∼10M duplicated reads per sample.

### In vitro footprinting

KLF2 (OriGene, TP310042) and RUNX2 (OriGene,TP760214) were incubated both separately and together with 25ng of naked BAC DNA (RP11-82D16) in a 22.5 uL volume of water and tagmentation buffer (20 mM Tris, 10 mM MgCl2 and 20% dimethylformamide). Incubation was carried out for 1 hour on a rotator at room temperature. 0.1 uL of preassembled Tn5 (seqWell, Tagify) in 2.4 ul of dilution buffer (50 mM Tris, 100 mM NaCl, 0.1 mM EDTA, 1 mM DTT, 0.1% NP-40 and 50% glycerol) was added to each reaction. Reactions (final TF concentration of 300 nM) were then tagmented at 37C for 30 minutes. Following tagmentation, DNA was purified with Qiagen MinElute columns, amplified for 8 cycles, and then purified once more with Qiagen MinElute columns prior to sequencing.

## Computational Methods

### Data Processing

All single-cell data have been processed according to the IGVF consortium standards and can be found: https://github.com/IGVF/single-cell-pipeline. Raw data, intermediate files, and code and workflows corresponding to preliminary analysis can be found on: https://data.igvf.org/analysis-sets/IGVFDS4798PRUI/

### TF Barcode Calling

Star (v2.7.10a) was used to generate a custom reference genome corresponding to the MORF library and to align FASTQ files. Following alignment, each sublibrary was processed independently to account for differences in sequencing depth. BAM files were filtered for alignment score, unique mapping, and polyG content (AS > 80, NH = 1, no polyG UMIs). Unique molecules (cell barcode + UMI + alignment) supported by 3 or more PCR duplicates were retained. TF enrichment per cell was assessed using a one-sided hypergeometric test, where the probability cell *i* was enriched for TF *j*:

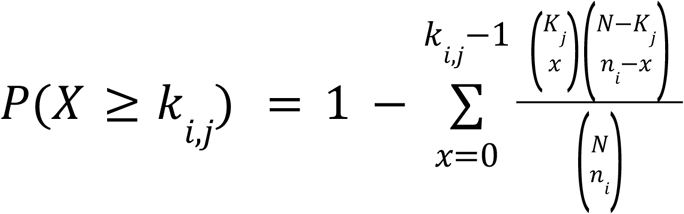

Where:

*k*_*i,j*_= TF *j* UMIs in Cell *i*

*K*_*j*_ = Sum of TF *j* UMIs in all cells

*N* = Sum of all UMIs in all cells

*n*_*i*_ = Sum of all UMIs in cell *i*

Hypergeometric testing was repeated using raw read counts. P values were adjusted using the Benjamini–Hochberg correction. TFs were retained if the adjusted *P* was < 0.05 for both UMI- and read-based tests.

### Protein structure modeling and evolutionary conservation

AlphaFold 3 was used with full length canonical amino acid sequences from Uniprot and the following DNA sequence and its reverse complement: 5’-GCCACAAACACGCCCACG-3’. Evolutionary conservation was determined using ConSurf^66^, under default parameters in sequence-mode.

### Robust topic modeling and single-cell topic rescoring

For both scRNA-seq and scATAC-seq analyses, topic models were fit separately within each modality using latent Dirichlet allocation (LDA) implemented in MALLET. To recover programs that are robust to noise in the data, we performed bootstrap resampling (10 replicates). For each bootstrap replicate, cells were sampled with replacement to a size of 500,000 cells. Each replicate was fit with 100 topics for 500 iterations using different random seeds.

To derive robust programs, topics recovered across bootstrap replicates were pooled, and pairwise topic similarity was computed from their topic x gene (RNA) or topic x peak (ATAC) feature loading vectors using jaccard similarity of the binarized matrices. A topic similarity graph was then constructed for visualization (Fig. 2B), and clustered using jaccard similarity to identify recurrent programs. Normalized gene or peak weights were averaged across topics within each recurrent cluster to obtain consensus RNA or ATAC topics.

Notably, due to the nature of bootstrapping, the original cell loadings for these topics only cover the subset of cells that were sampled. To obtain the consensus program activity scores for all cells, we adapted the chromVAR^67^ framework for both RNA and ATAC topics. For RNA, the cell x gene count matrix was used as input, and the motif x peak annotation matrix was replaced by the topic x gene matrix derived from the aforementioned process. Background genes were generated by sampling genes with matched average expression levels. For ATAC, topic activity was quantified analogously using the cell x peak accessibility matrix and the topic x peak matrix in place of motif annotations, with matched background peaks selected using the standard chromVAR background procedure. The resulting per-cell deviation scores were used as bias-corrected topic activity for visualization, cluster averaging, and downstream comparisons.

### Bulk seq2PRINT training, sequence attribution, and de novo motif discovery

scATAC-seq fragments from all cells retained for the final analysis were used to train the seq2PRINT model using the scPrinter package. Briefly, seq2PRINT is a sequence-to-footprint model that takes DNA sequences centered on candidate cis-regulatory elements (cCREs), together with their flanking regions (±920 bp), as input and predicts multi-scale chromatin accessibility footprints derived from the PRINT framework.

Sequence attribution scores were then computed across the peak set using DeepLIFT^68^, representing the contribution of each nucleotide in the input sequence to the predicted footprint and accessibility signals. De novo motifs were identified from these attribution scores using the seq_denovo_seq2print function in scPrinter, which adapts a TF-MoDISco-based clustering procedure to group high-contribution sequence segments. To better resolve composite motifs with different configurations, the default Leiden clustering partition (ModularityVertexPartition) was replaced with RBConfigurationVertexPartition with resolution set to 20. The discovered de novo motif matches within peaks were subsequently identified using finemo, which takes the de novo motifs and the corresponding seq2PRINT attribution scores as input for motif matching. Finemo hits were then summarized into peak x motif matrices for downstream analyses. De novo motifs were manually annotated when possible, indicated in Extended Table 3, with corresponding motif sequence logos in Extended Table 4.

### Patient Data Scoring

RNA-seq data and metadata from Fraietta et al.^42^ was downloaded. “Supplemental Table 5a: Transcriptomic profiling of mock-stimulated (control) CTL019 infusion products” was used for the RNA-seq data set. ATAC programs were connected to genes using DORC analysis^6^, retaining peak-to-gene associations with FDR < 0.05. After removing peaks with no gene association, the top 200 genes were retained, ordered by their associated peak’s ATAC program score. For RNA programs, the top 200 genes were selected, ordered by the RNA program score. Seurat 5.3.0 standard analysis was performed, and Seurat gene scoring was used on the 200 genes for each program for each patient sample. As in Fraietta et al., patients marked as complete remission (“CR”) and “PRTD” were grouped to indicate a positive response to therapy, and patients marked as partially responding (“PR”) and nonresponding (“NR”) were grouped to indicate a negative response to therapy.

### In vitro footprinting

Multiscale footprint scores for in vitro footprinting samples were calculated as in Nagaraja et al.^57^ Briefly, all conditions were downsampled to an equal number of total insertions; scores were then calculated with a one-sided binomial test for depletion of Tn5 insertions in center vs flanking windows of varying radii (2-100). To normalize for Tn5 bias, expected center vs flank distributions were calculated from naked DNA tagmentation controls. Footprint scores represent -log10(*P*) of depletion.

## Supplemental Oligo Table

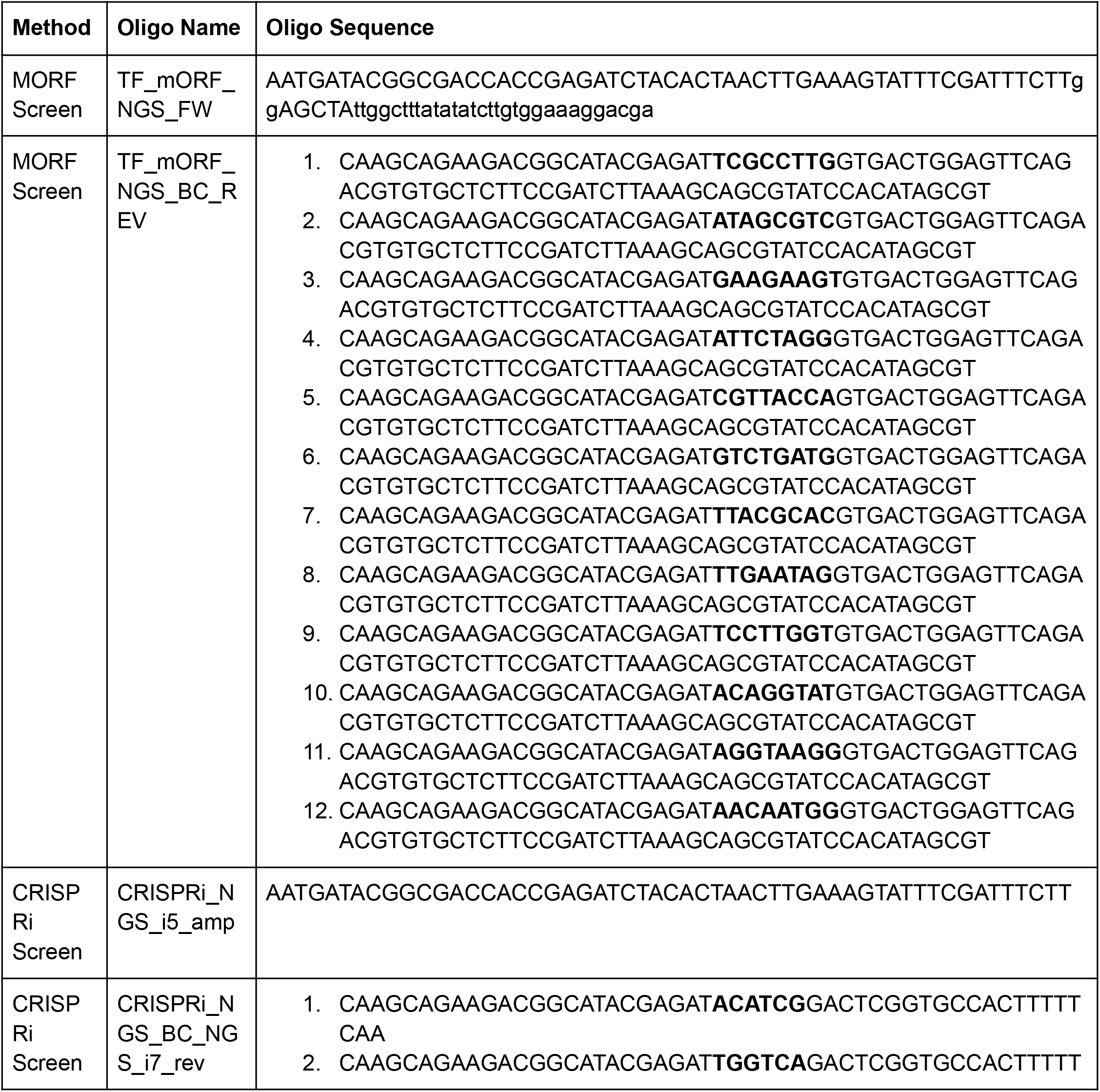
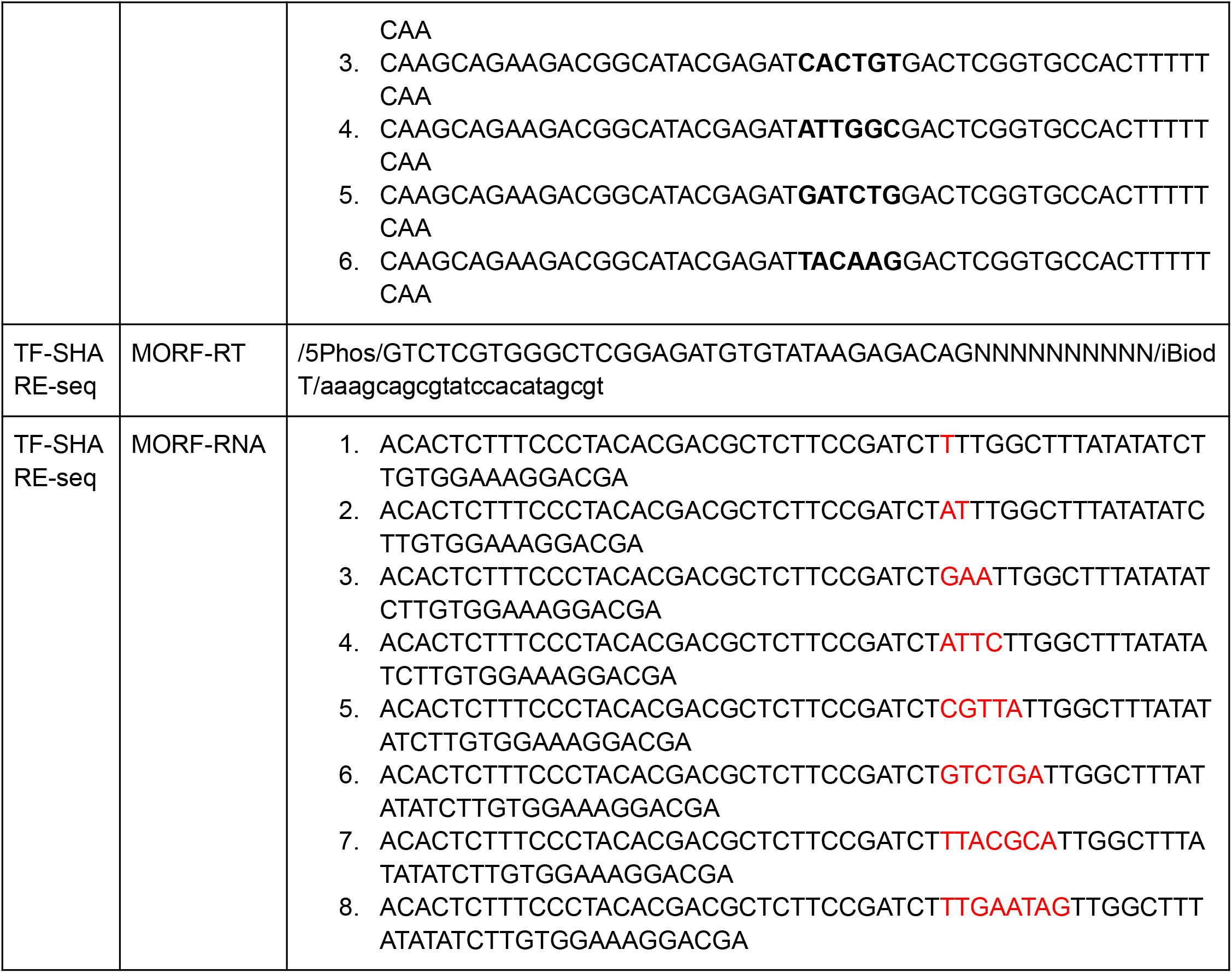

## Extended Tables

**Extended Table 1. MORF library information and screen results**. File contains the DESeq results (baseMean, log2FoldChange, lfcSE, stat, pvalue, and padj) for both the expanded (_Expanded) and exhausted (_Exhausted) screens. This file also contains whether the TF was profiled in perturb-SHARE-seq (“perturb_SHARE_seq_profiled” yes/no), and if it was, what shortened name was used for the TF in the manuscript (“perturb_share_name”). Additionally, this file contains isoform information from the original Joung et al. supplemental information, including: Source (source of the plasmid: Broad GPP or Genewiz), Name (TFORFxxxx), RefSeq_Gene_Name, RefSeq_and_Gencode_ID, Insert (DNA sequence), ORF_sequence (DNA sequence), Barcode_sequence (DNA sequence), and Epitope_Tag (None or V5).

**Extended Table 2. TF-Condition-Program Results**. File contains the program scores grouped by each TF in each condition. Specifically, it contains: TFOE (transcription factor overexpression ID), Condition (expanded: a.acute_unstim, expanded stim: b.acute_stim, exhausted: c.exhausted_unstim, exhausted stim: d.exhausted_stim), Topic (ATAC programs A1-21, RNA programs R1-17), Mean_Topic_Score (mean topic score across single cells, grouped by TFOE and Condition), SEM_Topic_Score (standard error of the mean of topic scores across single cells, grouped by TFOE and Condition), Mean_Topic_Score_Condition_Centroid (mean topic score across single cells, grouped by Condition), and SEM_Topic_Score_Condition_Centroid (standard error of the mean across single cells, grouped by Condition).

**Extended Table 3. De novo motif ID and naming**. File contains manual annotation of seq2PRINT de novo motifs. Specifically, it contains: DNM_id (a unique identifier of the de novo motif), seq2PRINT_id (the corresponding ID contained with Extended Table 4), Motif_Type (Footprint/Count based motif), Motif_Name (simplified name from the DNM_id), Composite_Status (Monomer/Composite/Unknown), Dimer_status (if it is a composite, if it is Homodimer/heterodimer/unknown).

**Extended Table 4. De novo motif logos**. File contains de novo motif logos. Specifically, it contains: pattern (the seq2PRINT_id), num_seqlets (the number of finemo seqlets that were merged), modisco_cwm_fwd (the logo in the forward orientation), modisco_cwm_rev (the motif logo in the reverse orientation), delta_effects (the aggregated footprint score surrounding the motif), and three putative computational matrices, given by name, q-value, and logo.

## Extended Data Figures

**Extended Data Figure 1.**
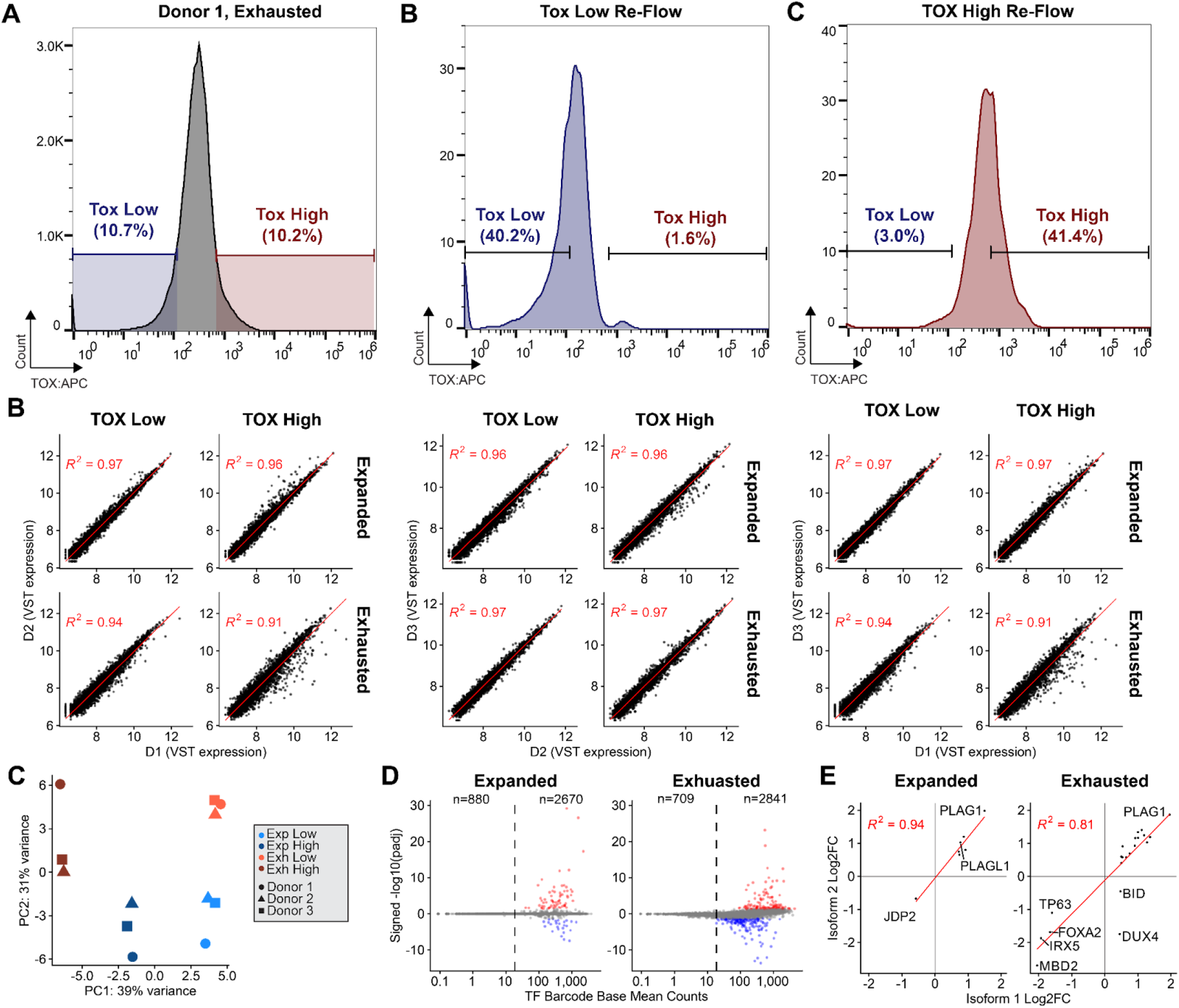
Quality control and validation of the TOX overexpression screen in primary human CD8+ T cells. (A) Representative flow cytometry histogram of TOX protein expression following exhaustion, with gating strategy used to re-sort TOX^HIGH^ and TOX^LOW^ populations. (B) Pairwise scatterplots of TF expression (variance-stabilized counts, VST) across donors for TOX^HIGH^ and TOX^LOW^ populations under expanded and exhausted conditions. Pearson R^2^ values are indicated in red. (C) Principal component analysis (PCA) of all samples, demonstrating clustering by condition and donor. (D) Mean TF barcode counts plotted against signed –log10 adjusted P values. Significantly enriched or depleted TFs (adjusted *P* < 0.05) are highlighted in red and blue, respectively. The dashed line indicates a base mean count of 18, below which statistical power was insufficient to reliably detect differential effects (E) Log2 fold changes (TOX^HIGH^ vs TOX^LOW^) for TF isoforms with at least two isoforms showing statistically significant effects.

**Extended Data Figure 2.**
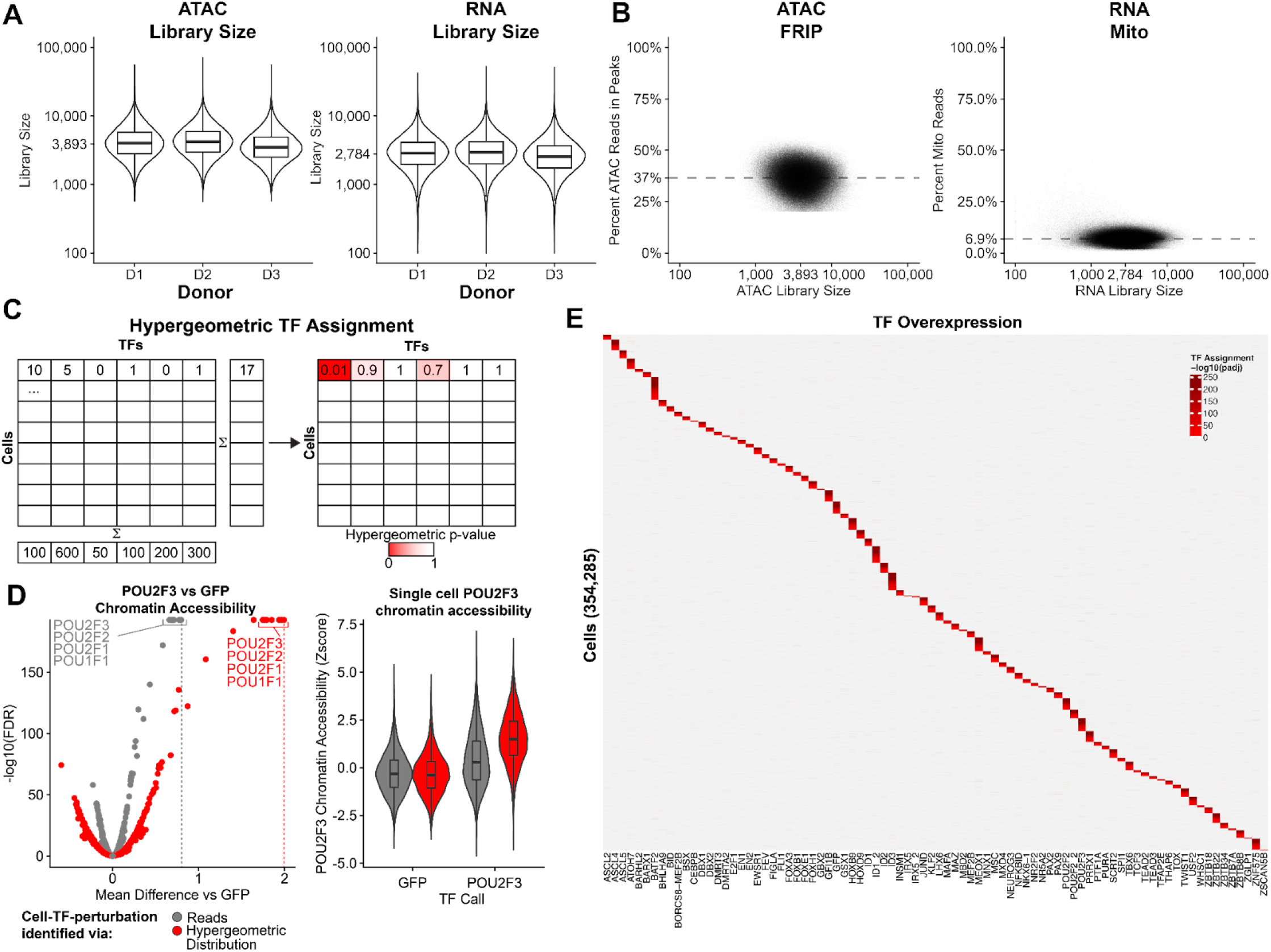
Perturb-SHARE-seq library quality. (A) ATAC-seq and RNA-seq library size across three biological donors. (B) ATAC fraction of reads in peaks (FRIP) as a function of library size (left); dashed line indicates the mean FRIP (37%). RNA mitochondrial read percentage as a function of library size (right); dashed line indicates the mean mitochondrial content (6.9%). (C) Schematic illustrating hypergeometric assignment of transcription factor (TF) barcodes to single cells. (D) Left, Welch’s two-sample t-test comparing motif accessibility between POU2F3- and GFP-expressing cells, defined by either detection of TF barcode reads (grey) or significant hypergeometric enrichment (red). Right, violin plots showing POU2F3 motif accessibility at single-cell resolution. (E) Heatmap of TF assignment probabilities across single cells; grey denotes adjusted P ≥ 0.05 and red denotes adjusted *P* < 0.05.

**Extended Data Figure 3.**
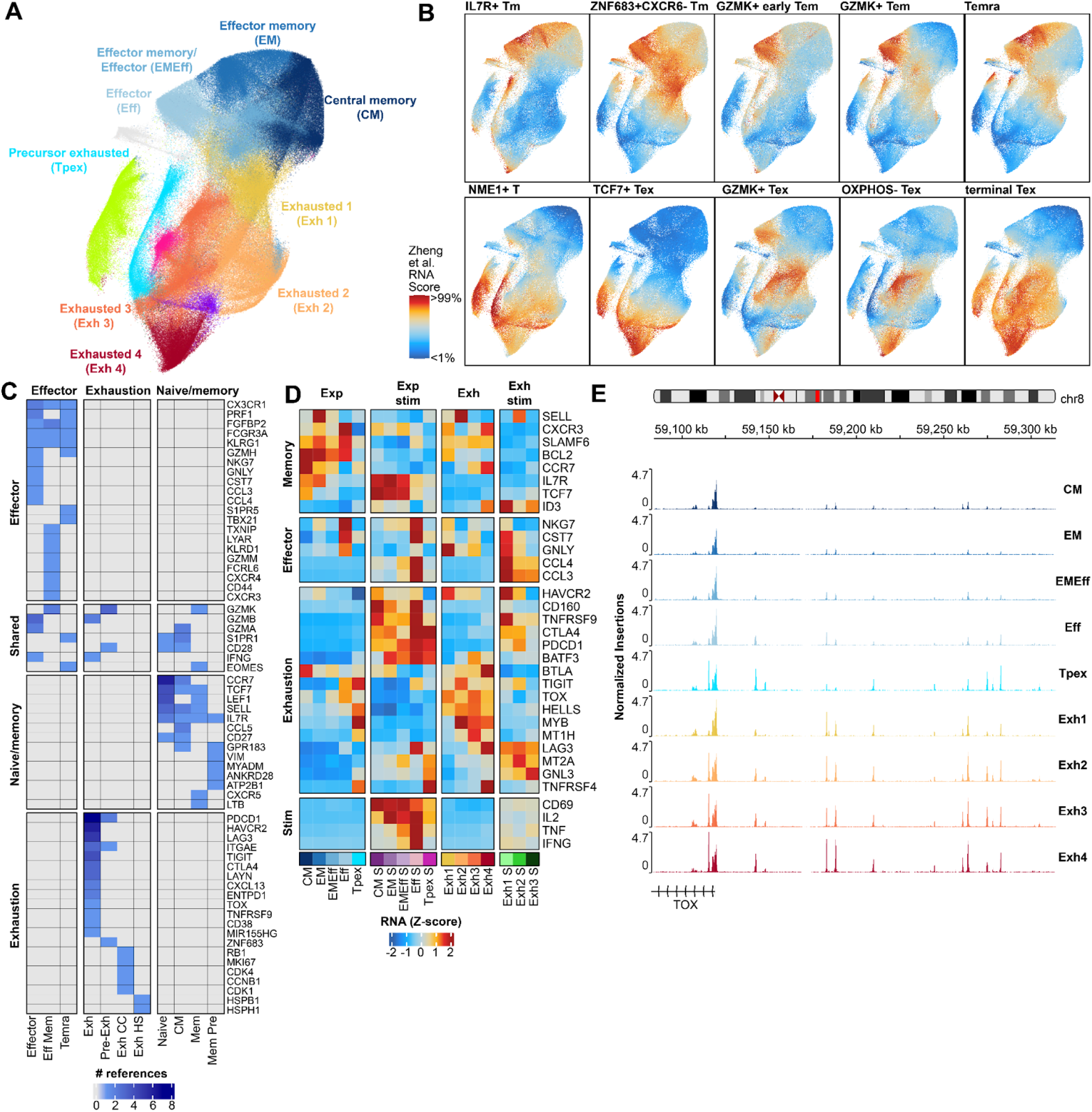
Annotation of CD8+ T cell states and associated chromatin accessibility. (A) UMAP of unstimulated cells based on cisTopic embedding, colored by Louvain clusters. Synthetic clusters are unlabeled (green, OCT TFs; pink, bHLH TFs; purple, HOX TFs; grey, cluster 19). (B) UMAP of unstimulated cells colored by Zheng et al. RNA signature scores for CD8+ T cell states. (C) Summary of literature-derived marker genes for CD8+ T cell states, colored by the number of supporting references linking each gene to a given state. (D) Heatmap showing z-scored mean RNA expression of CD8+ T cell marker genes (rows) across Louvain clusters (columns). Exp: expanded, Exh: exhausted (E) Genome browser tracks showing chromatin accessibility at the TOX locus across unstimulated CD8^+^ states.

**Extended Data Figure 4.**
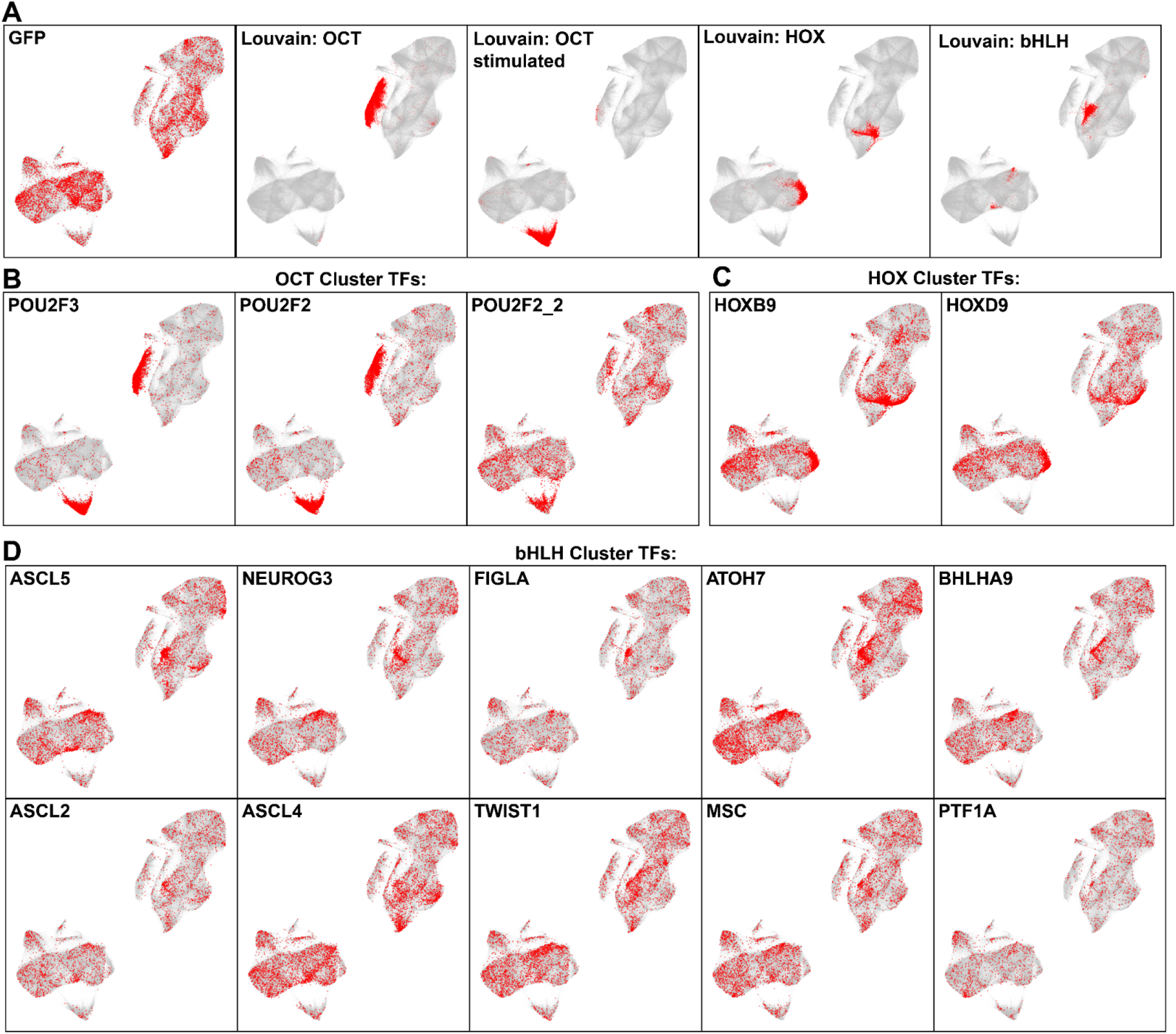
Synthetic clusters defined by transcription factor overexpression. (A) UMAP embedding of all cells (grey), with cells overexpressing GFP highlighted in red. Cells are additionally grouped by Louvain clustering, with cluster identities indicated in the title. (B) UMAP of all cells (grey), with cells overexpressing OCT family transcription factors highlighted in red. (C) UMAP of all cells (grey), with cells overexpressing HOX family transcription factors highlighted in red. (D) UMAP of all cells (grey), with cells overexpressing selected bHLH transcription factors highlighted in red.

**Extended Data Figure 5.**
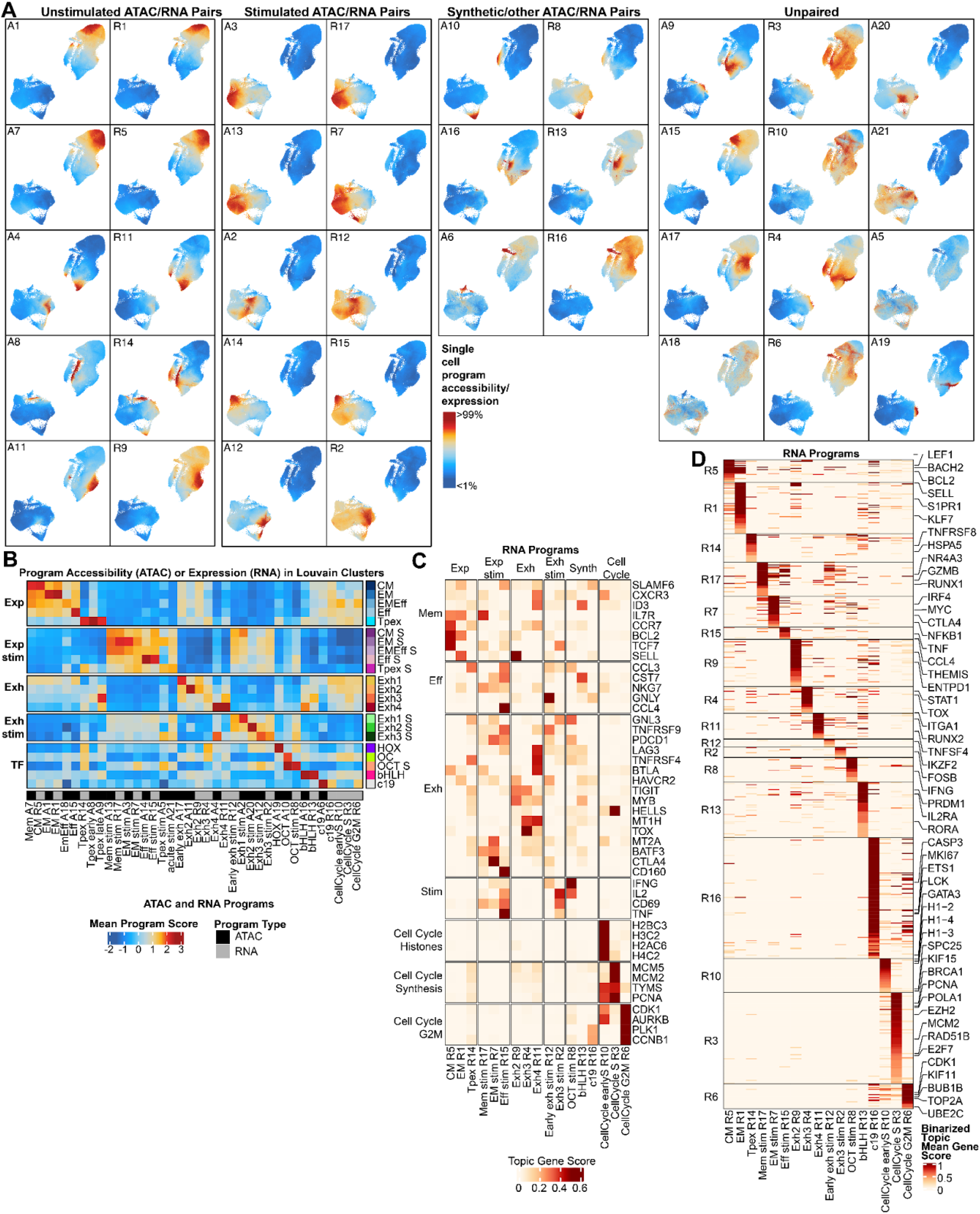
Characterization of ATAC and RNA programs. (A) UMAP embeddings colored by single cell chromatin program accessibility (A) or RNA expression (R). Programs are grouped into unstimulated ATAC-RNA pairs, stimulated ATAC-RNA pairs, synthetic and other pairs, and unpaired programs. Color scale ranges from the 1st to 99th percentile. (B) Heatmap of ATAC program accessibility and RNA program expression, shown as mean values across single cells and grouped by cluster identity. (C) Heatmap of RNA program gene scores for selected marker genes, showing that programs cluster according to marker gene expression patterns. (D) Heatmap of binarized mean RNA gene scores, highlighting the top marker genes for each RNA program.

**Extended Data Figure 6.**
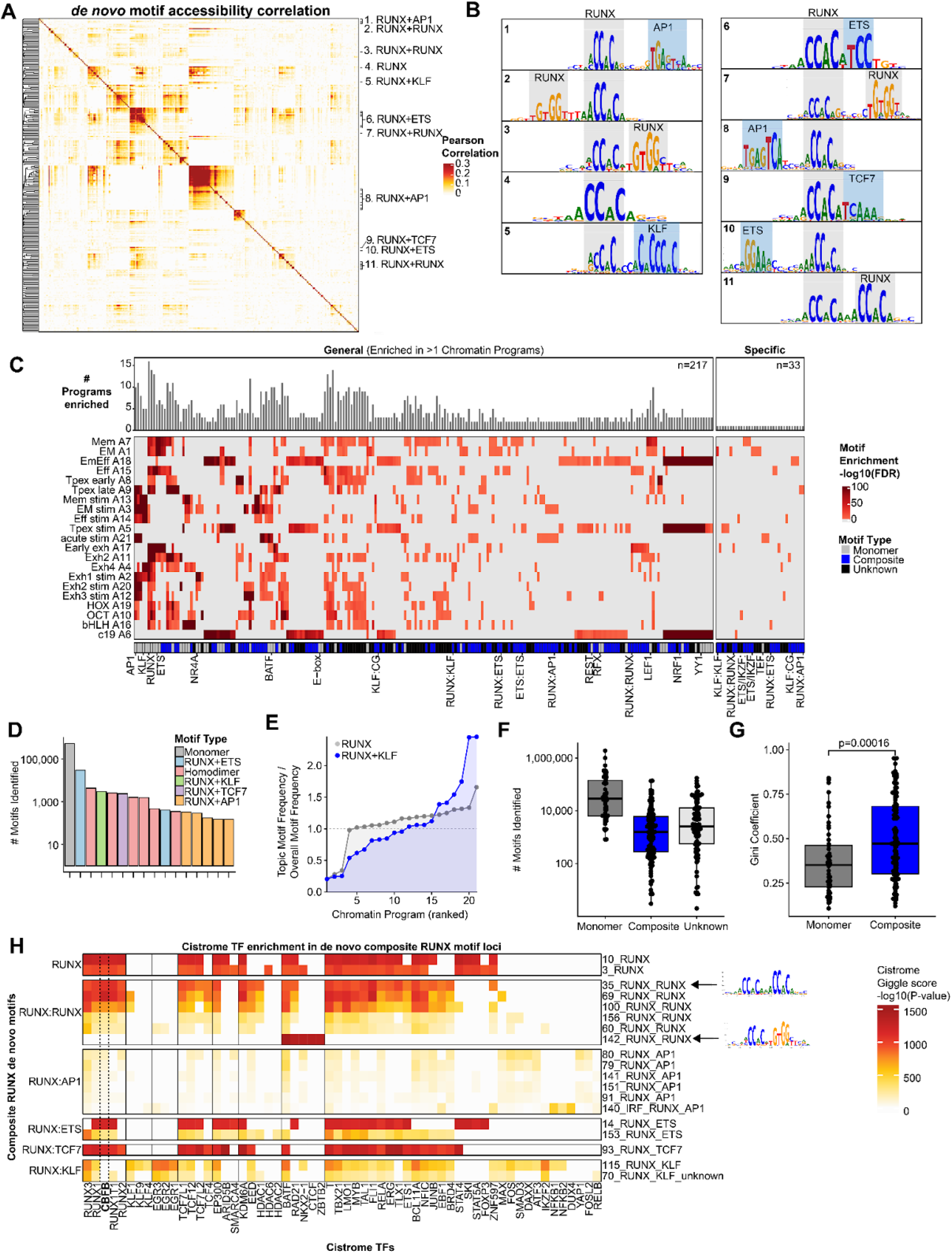
De novo motif discovery using seq2print reveals monomeric and composite motif architecture. (A) Pearson correlation of de novo motif accessibility across single cells. RUNX monomeric and composite motifs are highlighted on the right. (B) Representative sequence logos of highlighted RUNX-containing de novo motifs in (A). (C) Heatmap of de novo motif enrichment (columns) across chromatin programs (rows), colored by -log10 hypergeometric FDR. Motifs are annotated by type (grey, monomeric; blue, composite; black, unknown). Top barplot shows the frequency of significantly enriched topics for each de novo motif. (D) Bar plot showing the number of motifs identified within RUNX motif subsets, colored by monomer and composite types. (E) Scatter plot of motif frequency within topics versus overall motif frequency. Each point represents a topic; the y-axis shows enrichment (topic motif frequency / overall motif frequency). Blue, RUNX–KLF composite motifs; grey, RUNX monomeric motifs (F) Box plot of the total motifs identified for each de novo motif, across monomer and composite motifs. (G) Box plot of Gini coefficients of motif enrichment across topics, split by monomer and composite motifs. (H) Heatmap of cistrome enrichment of TF colocalization with de novo motif genomic loci, colored by GIGGLE score (-log10 *P*). CBFB indicated with a dashed line.

**Extended Data Figure 7.**
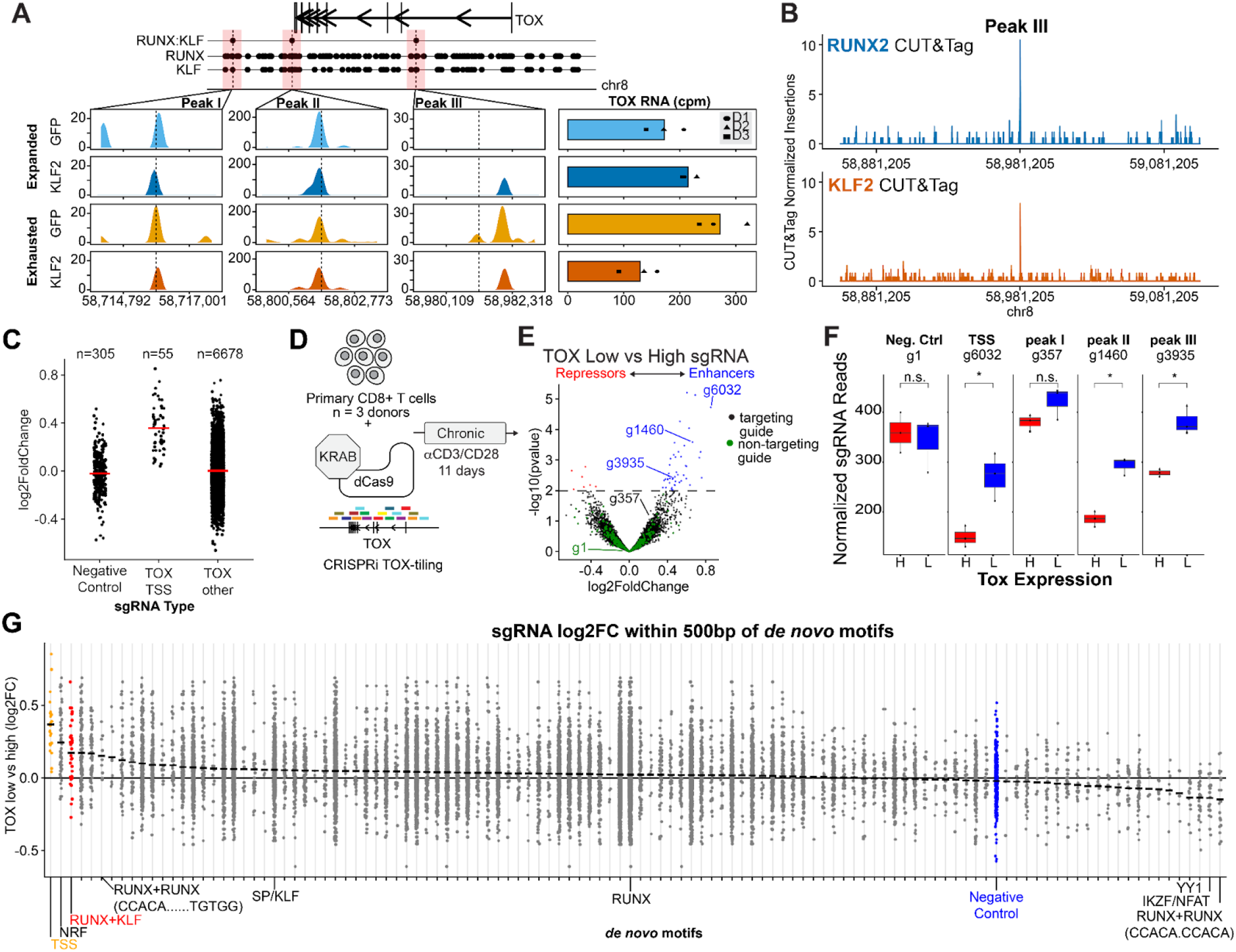
CUT&Tag and CRISPRi dissection of the TOX locus. (A) Top: Genomic locations of de novo motifs corresponding to RUNX:KLF composite sites, as well as RUNX and KLF monomer motifs within the TOX locus. Regions containing RUNX:KLF composite motifs are labeled as peaks I–III. Bottom: Pseudobulk chromatin accessibility tracks at the three composite peaks in GFP and KLF2-overexpressing cells under expanded and exhausted conditions (left). Dashed lines mark the positions of RUNX:KLF composite motifs. TOX RNA expression in GFP and KLF2 pseudobulks, stratified by donor (right). (B) CUT&Tag profiles of RUNX2 and KLF2 binding at peak III, normalized to fragments per million. (C) CRISPRi sgRNA log2 fold change (TOX-low vs -high), stratified by sgRNA class: negative controls, guides targeting the TOX transcription start site (TSS), and all other guides across the locus. Red lines indicate the mean log2 fold change for each group. (D) Schematic of CRISPRi guide tiling across regulatory elements at the TOX locus. (E) Volcano plot of CRISPRi guide enrichment in the TOX high vs TOX low populations. Guides decreasing TOX expression (blue), increasing TOX expression (red), and negative controls (green) are highlighted. (F) Representative CRISPRi guides across three donors, showing read distribution in TOX high (red) and TOX low (blue) bins. (G) sgRNA log2 fold change for guides located within 500 bp of de novo motifs, grouped by motif. Black horizontal bars indicate the mean log2 fold change per motif. Motifs are ordered by mean effect size.

**Extended Data Figure 8.**
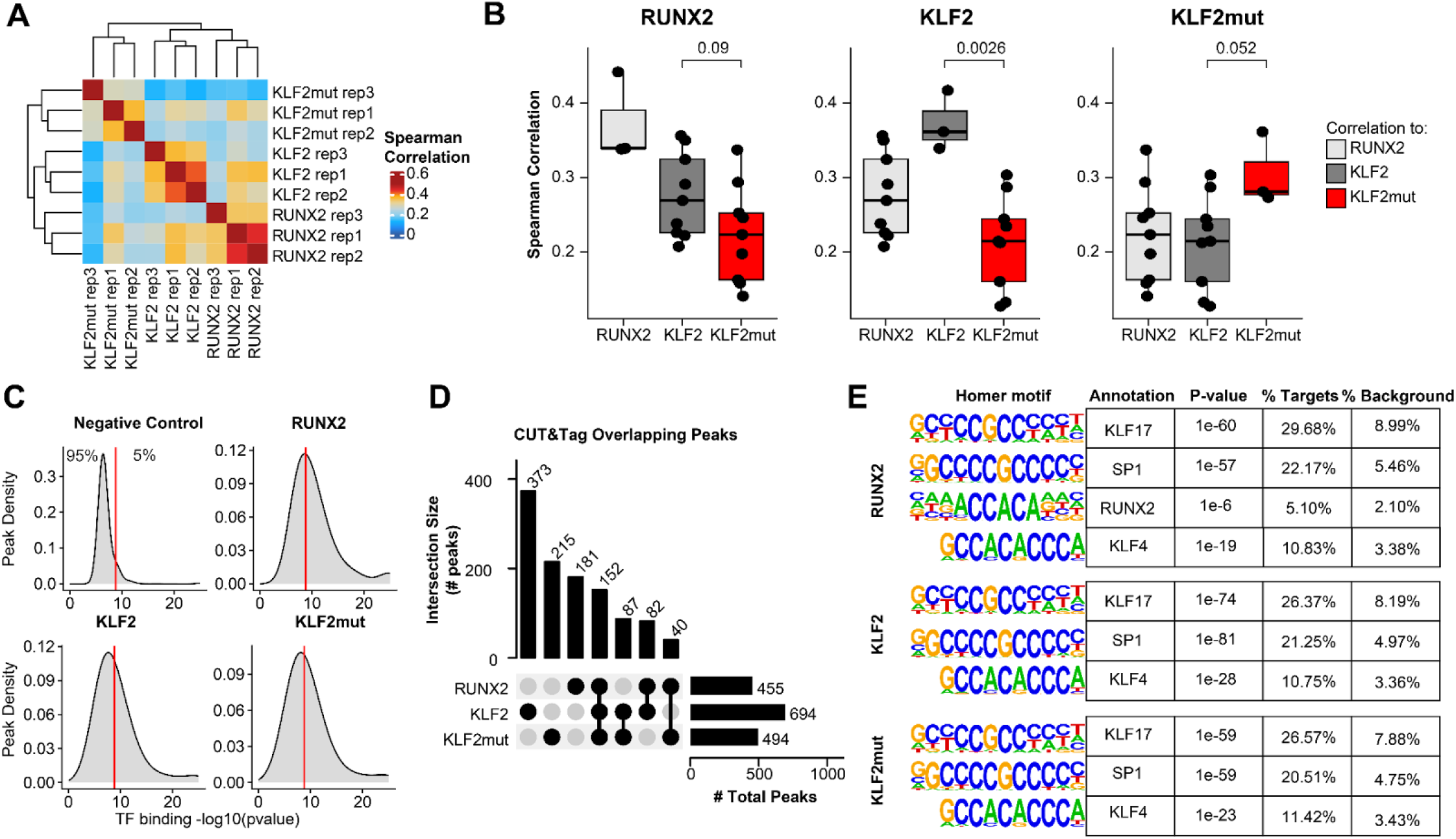
CUT&Tag profiling of RUNX2 and KLF2 variants reveals reproducible binding and motif enrichment. (A) Spearman correlation of k-mer accessibility across RUNX2, KLF2, and KLF2mut CUT&Tag samples, with hierarchical clustering. (C) Box plots of Spearman correlation across samples, excluding self-correlations (same TF, same replicate). (D) Density plot of MACS2 peak scores (-log10 P value) for peaks identified in negative control (no primary antibody), RUNX2, KLF2, and KLF2mut samples. The red line indicates the threshold at which 5% of negative control peaks are retained. (E) UpSet plot showing overlap of peaks (passing the threshold in C) across samples. Intersection sizes are shown above, and total peak set sizes are shown on the right. (F) HOMER motif logos of known motifs identified from RUNX2, KLF2, and KLF2mut CUT&Tag peaks.

**Extended Data Figure 9.**
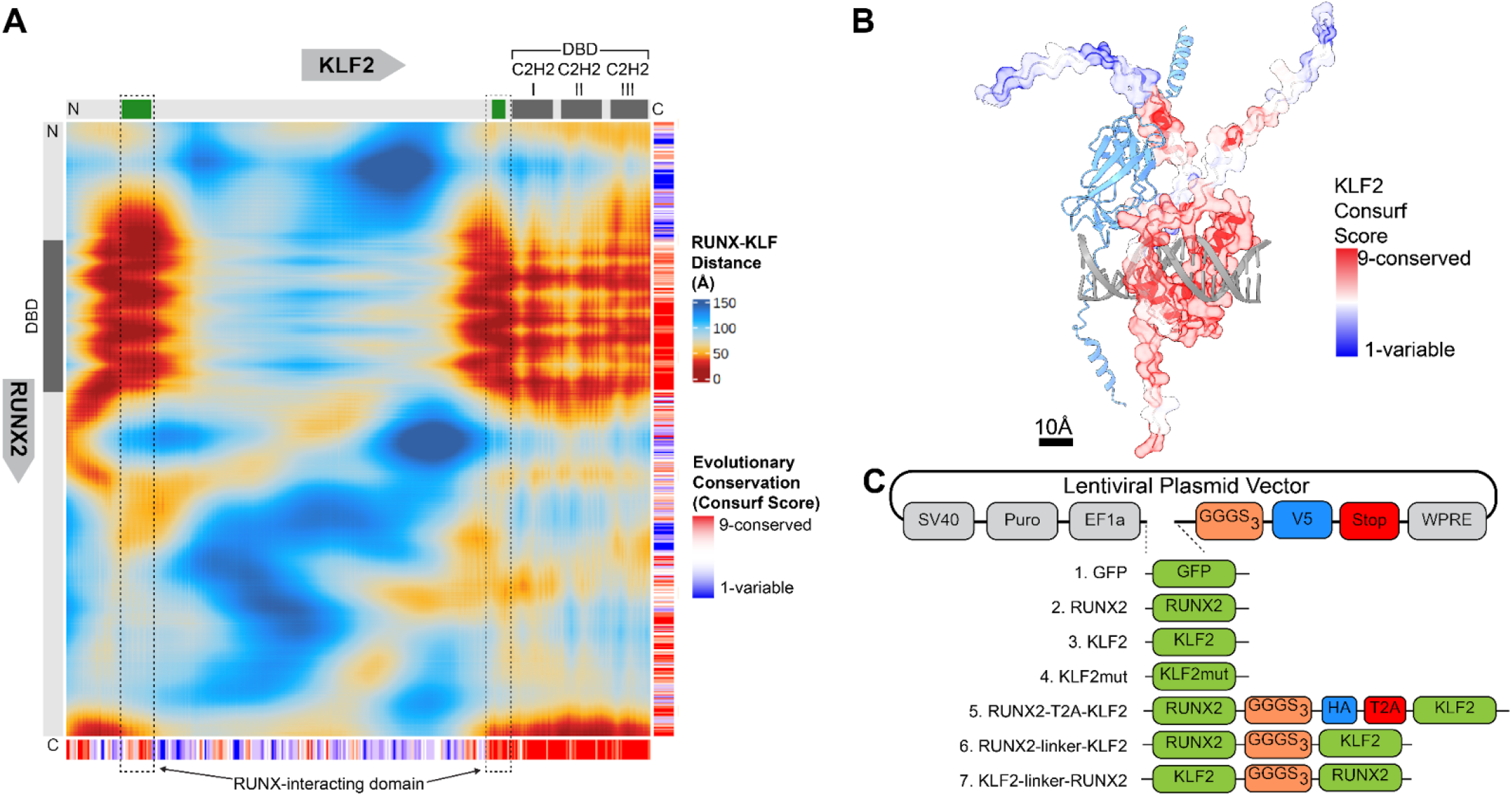
RUNX2–KLF2 structural analysis and construct design. (A) Heatmap showing proximity between each amino acid in KLF2 (x-axis; N- to C-terminus, left to right) and RUNX2 (y-axis; N- to C-terminus, top to bottom), colored by minimum heavy-atom distance (Å). Conservation scores (ConSurf) are shown for KLF2 (bottom) and RUNX2 (right), with low conservation in blue and high conservation in red. Protein structural features, including DNA-binding domains, are annotated along the axes. (B) AlphaFold3-predicted structure of RUNX2 (blue) in complex with DNA and KLF2, with KLF2 colored by ConSurf conservation score. (C) Schematic of plasmid constructs used for transcription factor overexpression experiments. TFs are shown in green, linkers in orange, HA and V5 tags in blue, and stop codons and T2A sequences in red.

